# Cell type-specific roles of FOXP1 in the excitatory neuronal lineage during early neocortical murine development

**DOI:** 10.1101/2024.06.08.598089

**Authors:** Ana Ortiz, Fatma Ayhan, Nitin Khandelwal, Elliot Outland, Miranda Jankovic, Matthew Harper, Genevieve Konopka

**Author notes:** **Lead Contact information** Genevieve Konopka, Ph.D. Department of Neuroscience, University of Texas Southwestern Medical Center, 5323 Harry Hines Blvd., ND4.300, Dallas, TX 75390-9111.

## Abstract

FOXP1, a transcription factor enriched in the neocortex, is associated with autism spectrum disorders (ASD) and FOXP1 syndrome. *Emx1^Cre/+^;Foxp1^fl/fl^*conditional deletion (*Foxp1* cKO) in the mouse cortex leads to overall reduced cortex thickness, alterations in cortical lamination, and changes in the relative thickness of cortical layers. However, the developmental and cell type-specific mechanisms underlying these changes remained unclear. We find that *Foxp1* deletion results in accelerated pseudo-age during early neurogenesis, increased cell cycle exit during late neurogenesis, altered gene expression and chromatin accessibility, and selective migration deficits in a subset of upper-layer neurons. These data explain the postnatal differences observed in cortical layers and relative cortical thickness. We also highlight genes regulated by FOXP1 and their enrichment with high-confidence ASD or synaptic genes. Together, these results underscore a network of neurodevelopmental disorder-related genes that may serve as potential modulatory targets for postnatal modification relevant to ASD and FOXP1 syndrome.

## Introduction

The neocortex is radially organized into layers, each of which is enriched with specialized subtypes of neurons generated from deepest to superficial layers to form its laminar structure. Excitatory glutamatergic neurons are sequentially generated from progenitors residing in two germinal compartments — the ventricular zone (VZ) and the subventricular zone (SVZ) — lining the cerebral ventricles. Apical radial glia (aRGs) in the VZ divide to either self-renew or directly generate neurons ^1,2^. aRGs also indirectly generate neurons by producing intermediate progenitor cells (IPs) whose cell bodies reside in the SVZ ^3^. These IPs further self-amplify and therein boost neuronal production ^4-6^. In mice, the generation of excitatory neurons for a specific cortical layer peaks at different embryonic days in an inside-out manner. Deep layer neurons are born first (L5-6), followed by the generation of neurons for the upper layers (L2-4) ^7-13^.

Development of the neocortex requires the orchestrated execution of a series of crucial processes. Deviations in this fine-tuned process may increase susceptibility to neurodevelopmental disorders ^14^. One such disorder is FOXP1 syndrome that is caused by mutations or deletions in the *FOXP1* gene ^15-17^. FOXP1 (Forkhead Box Protein P1) is a transcription factor that plays a crucial role in various developmental processes, including cortical development. Individuals with FOXP1 syndrome often exhibit a range of neurological and behavioral symptoms, including intellectual disability, language impairments, motor delays, and autism spectrum disorders (ASD) ^18-20^. Moreover, many individuals exhibit ADHD symptoms, such as inattention and hyperactivity ^18,19^. However, the specific features of FOXP1 syndrome can vary widely ^21^. Previous work suggests that FOXP1 regulates other genes encoding proteins with roles in neural development ^22,23^. However, further insight into the cellular and molecular mechanisms underlying these changes in the cortex is needed.

Mouse models with *Foxp1* deletion have shown phenotypes relevant for individuals with FOXP1 syndrome. *Foxp1* whole-body heterozygous or brain-specific knock-out mice exhibit reduced neonatal ultrasonic vocalization (USVs), hyperactivity, and metabolic disturbances and mitochondrial dysfunction in striatal neurons ^24-27^. Further, knocking-down *Foxp1* in mice via *in utero* electroporation during mid-neurogenesis results in broad migration deficits observed at late embryonic stages ^23,28^. Additionally, using the same mice as in the current study, we previously reported that loss of FOXP1 in the dorsal forebrain of mice results in ASD-relevant behaviors including neonatal and adult USVs, social interaction deficits, hyperactivity, spatial learning deficits, and also impairs synaptic plasticity ^29,30^. Observed neuroanatomical changes include a reduction in neocortical size and abnormal positioning of neurons that express canonical markers of upper layer neurons in the deep layers of the postnatal mouse neocortex ^30^. However, the molecular changes underlying these phenotypes remained undefined.

Postnatally, FOXP1 exhibits layer-specific expression, detected in neurons of L3-5 and sparsely in L6a neurons ^30,31^. During embryonic development, FOXP1 also has restricted expression in distinct cell types across neocortical development, with expression in both proliferative cells and post-mitotic neurons during early neurogenesis and confined to post-mitotic neurons during late neurogenesis ^23,32^. In proliferative cells outside of the nervous system, FOXP1 acts as a transcriptional repressor ^33,34^. Research in embryonic stem cells shows that FOXP1 preferentially binds distinct sequence motifs compared to differentiated cells, thereby regulating the expression of genes associated with pluripotency versus differentiation ^35^. Together, these data raise the question of whether FOXP1 serves a distinct cell type-specific roles across neocortical development.

In this study, we sought to delineate the molecular contributions of FOXP1 in a cell type-specific manner across neocortical development. We investigated the gene regulatory network governed by FOXP1 during murine neocortical development, focusing on its cell type-specific functions. We generated *Foxp1 ^flox/flox^* conditional knockouts (*Foxp1* cKOs) by selectively deleting *Foxp1* from excitatory neocortical neural progenitors using the *Emx1*.Cre mouse line ^36^. Utilizing single-nucleus RNA sequencing (snRNA-seq) and single-nucleus Assay for Transposase-Accessible Chromatin sequencing (snATAC-seq), we ascertained the genetic and epigenetic regulatory mechanisms controlled by FOXP1 in different cell types at various ages. Additionally, we sought to investigate whether FOXP1 loss affects cell fate specification or migration.

We find that FOXP1 exerts opposing and congruent regulation on gene expression and chromatin accessibility during early neocortical development, particularly impacting genes associated with ASD and synaptic function. These are compelling mechanistic insights into our previous behavioral studies in these *Foxp1* cKOs that demonstrated ASD-relevant altered behaviors. We also report enrichment of FOX motifs in a cell type-specific manner, indicating genomic sites for binding and regulation by FOXP1. Using thymidine analogue pulsing, we showed a change in cell cycle exit rates in *Foxp1* cKOs during late neurogenesis. Additionally, thymidine analogue birth-dating revealed migration deficits in a subpopulation of upper-layer (UL) neurons and identified genetic players potentially contributing to this phenotype. Overall, FOXP1 loss affects (i) genes encoding proteins associated with ASD and synaptic function, (ii) cell cycle exit, (iii) cell type-specific transcription factors, (iv) cell type proportions, and (v) migration of specific UL neurons.

Understanding the dynamic genetic and epigenetic landscape during development is crucial for deciphering typical and atypical neural development. Our study elucidates the role of FOXP1 at key neurogenic stages and provides insights into potential convergent disease mechanisms underlying distinct molecular etiologies.

## Results

### snRNA-seq and snATAC-seq identified DEGs and DARs in cell types with excitatory neuronal lineage at E13.5, E16.5 and P0.5 time points

To determine the cell type-specific roles of FOXP1 across early neocortical development, we generated *Foxp1 cKO* and control littermates by crossing *Foxp1^flox/flox^* mice with mice expressing *Emx1.Cre* ^33,36^. This induces recombination embryonically in excitatory neurons and progenitors derived from the dorsal telencephalon. We dissected whole-cortex and used the 10X Genomics platform to perform snRNA-seq and snATAC-seq (Supplemental Fig. 1, 2).

As FOXP1 exhibits temporal regulation in distinct cell types throughout development in the neocortex, we selected three timepoints for sequencing: embryonic day (E) 13.5, E16.5, and postnatal day (P)0.5. At E13.5, FOXP1 is expressed in neuronal progenitors and post-mitotic neurons, but by E16.5 FOXP1 expression is enriched in neurons and sparsely expressed in neuronal progenitors (Supplemental Fig. 3) ^32^. Postnatally FOXP1 exhibits layer-restricted expression in L3-5 and sparse expression in L6a ^30-32^.

The snRNA-seq dataset of the whole cortex (“*all cells*” dataset) captures expected cell types found in a developing neocortex such as choroid plexus, microglia, and neuronal cells (Supplemental Fig. 1). The total nuclei per timepoint consists of E13.5: 56,557; E16.5: 36,708; P0.5: 84,075 (Supplemental Fig. 1). Cell types were annotated using canonical markers and existing neocortical datasets ^37,38^ (see Methods). However, the *Emx1.Cre* is known to recombine mostly in the progenitors of the excitatory neuronal lineage. When analyzing glial cells that arise from shared progenitors with excitatory neurons, we found limited changes (Supplemental Fig. 1D-E). Further, we previously reported cytoarchitectural changes in the postnatal brain in excitatory post-mitotic neurons, such as reduced excitatory neuron numbers and changes in cortical volume ^29,30^. Thus, we subset only cell types from the excitatory neuronal lineage to focus downstream analyses (“excitatory lineage” dataset, Fig. 1A,B).

**Figure 1:**
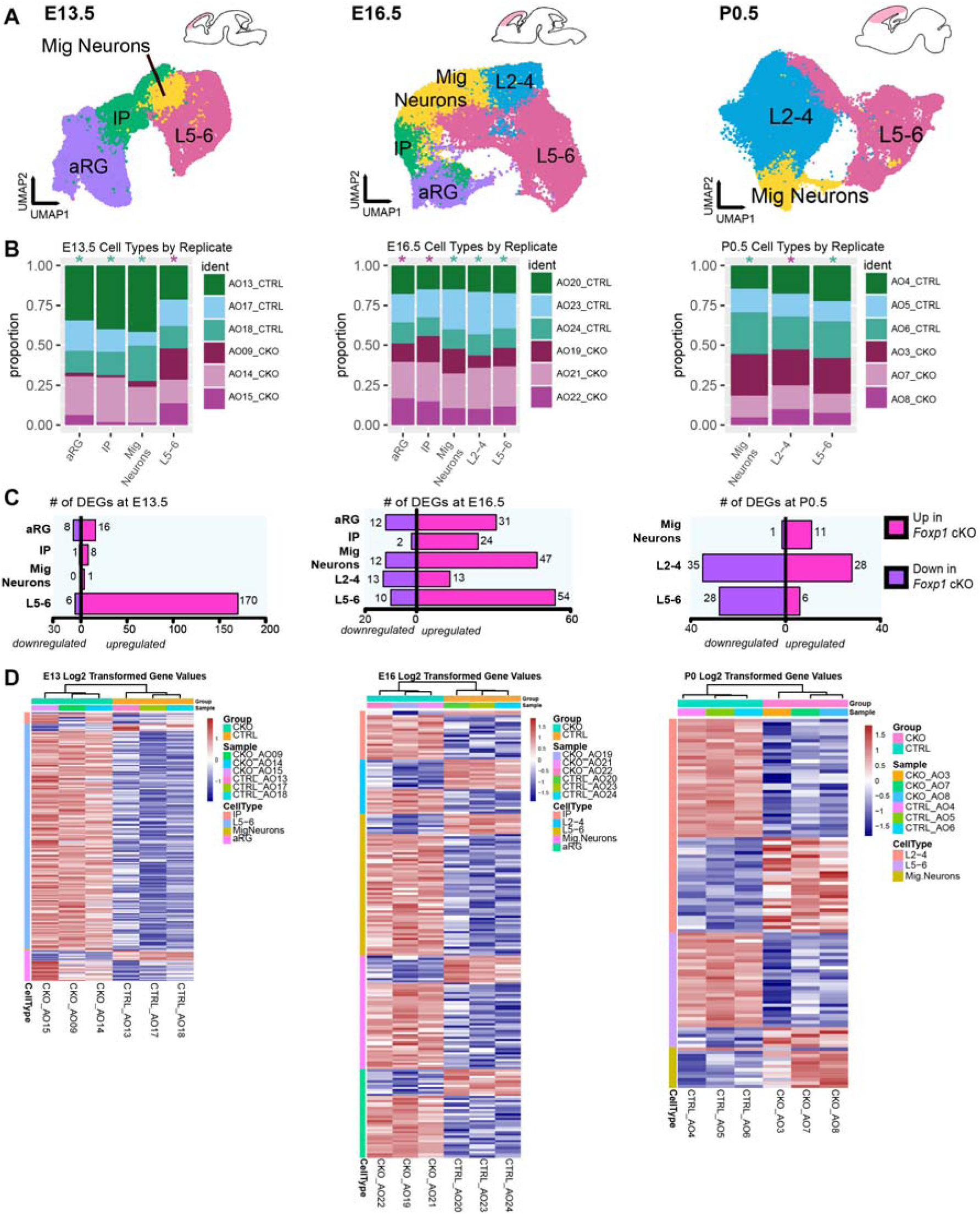
Excitatory neuronal lineage snRNA-seq of *Foxp1* cKO and control cortices shows disruptions in gene expression. **A**) Processed snRNA-seq data from all timepoints demonstrates expected cell types and cell distributions across neocortical development. Total cells per timepoint are E13.5: 26,513; E16.5: 12,011; P0.5: 43,040. N=3/genotype. **B)** Stacked bar plots show representation from replicates across broad cell type clusters. Star above each bar indicates a significant difference in cell type proportion between genotypes using the two-proportions z-test with Yates correction; green for control and purple for cKO. **C)** Number of DEGs per broad cell type by timepoint. DEGs were defined as significant with adj. p-val < 0.05 and average |logFC| > 0.15. **D**) Heatmaps showing the expression of DEGs for each cell type by biological replicate.

After subsetting and re-clustering, we detected expected cell types such as aRGs, IPs, migrating neurons, and neurons from L5-6 or L2-4, which are differentially colored by cell type in the UMAP representation (Fig. 1A, Supplemental Fig. 4). We observed 16 clusters at E13.5, 21 at E16.5, and 23 at P0.5 (Supplemental Fig. 4A-C). We collapsed cell type clusters into “broad” cell type clusters (*CellType_coll*) for downstream analyses and only show these clusters for ease of visualization (Fig. 1A). Across the three timepoints, broad cell type clusters had representation from all biological replicates (Fig. 1B). The composition of cell types also changes as expected across our selected timepoints (Supplemental Fig. 4D).

Analysis of gene expression differences showed that the *Foxp1* loss led to a notable increase in the number of upregulated differentially expressed genes (DEGs) in all cell types at embryonic timepoints and only in migrating neurons at P0.5 (Fig. 1C). FOXP1 is known to act as a transcriptional repressor, especially in cells with a proliferative capacity ^33,34^. Therefore, the observed pattern of more upregulated DEGs in *Foxp1* cKOs compared to controls supports a role for FOXP1 as a transcriptional repressor during early neocortical development. In contrast, at P0.5, L5-6 and L2-4 neurons demonstrated more downregulated DEGs, indicating a potential shift in the role of FOXP1 postnatally (Fig. 1C). The expression of the DEGs was relatively consistent among the biological replicates for each genotype at each timepoint (Fig. 1D). Thus, our stringent filtering and analysis methods resulted in a robust list of gene expression changes downstream of loss of FOXP1 in the embryonic cortex. Further, these DEGs show overlap with a P0 whole cortex bulk RNA-seq dataset we previously generated (Supplemental Fig. 4E-F). However, the overlap is highest at P0 in our snRNA-seq pseudo-bulk analysis, which makes sense given the matching sample ages.

We also performed snATAC-seq on whole cortical tissue (Supplemental Fig. 2) and annotated the data using label transfer from our snRNA-seq dataset (see Methods). We subset the excitatory neuronal lineage to facilitate comparative analysis with our excitatory lineage snRNA-seq data. Using broad cell-type annotation (*CellType_coll*), we identified all cell types that were detected in our snRNA-seq data, ensuring consistency across our datasets (Fig. 2A). Each cluster showed representation from each biological replicate (Fig. 2B). To understand the potential regulatory role of FOXP1 in chromatin accessibility, we identified differentially accessible regions (DARs) between genotypes (see Methods, Fig. 2C). In contrast to our observation of more upregulated DEGs across all embryonic timepoints and cell types, and migrating neurons at P0.5, there are more closed DARs at E13.5 and P0.5 but not at E16.5 (Fig. 2C). The distribution of DARs among genomic regions is relatively consistent across timepoints and open/closed status, with a notable exception of an increased proportion of DARs in the first intron of genes within the P0 closed group (Fig. 2D).

**Figure 2:**
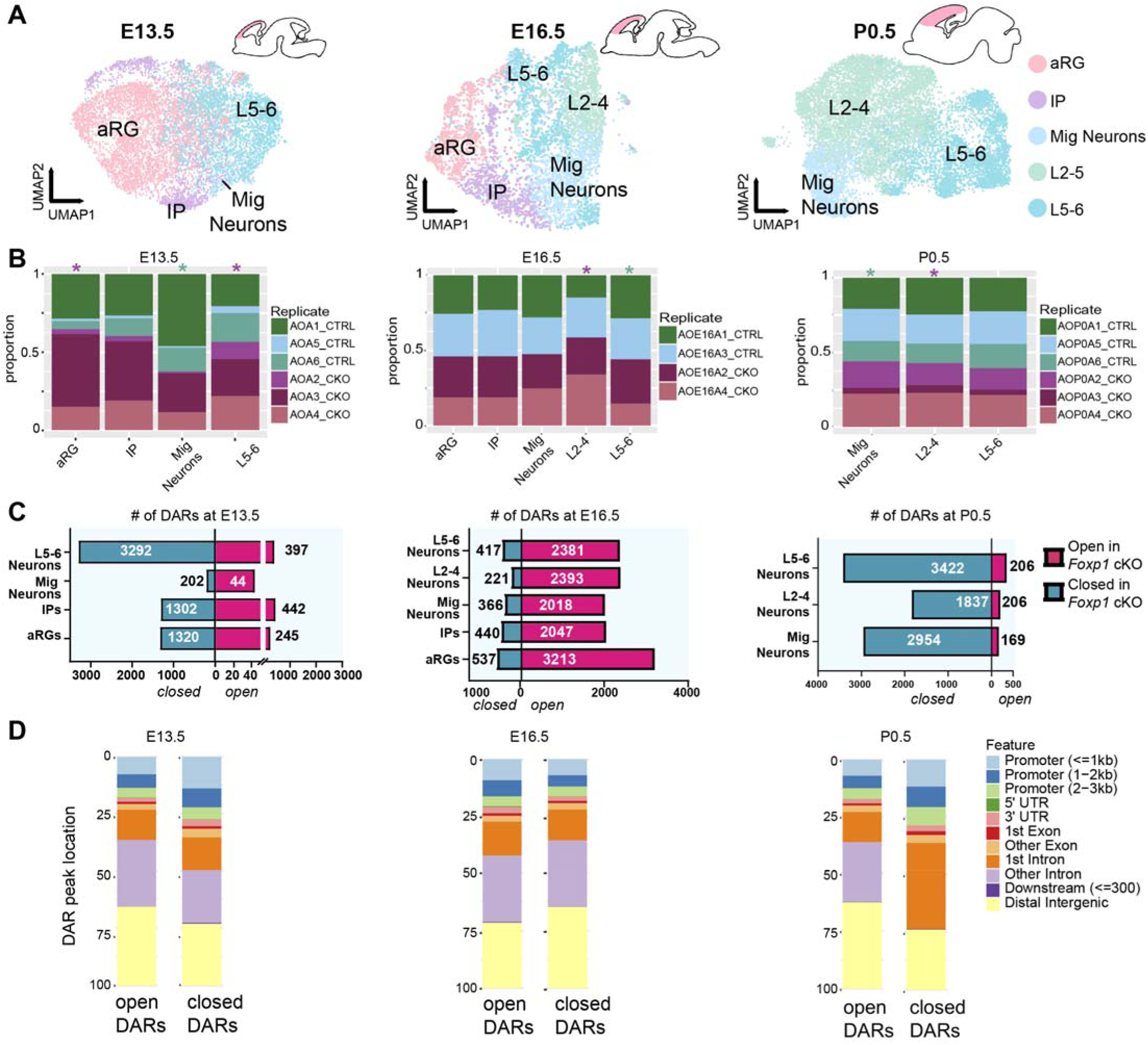
Excitatory neuronal lineage snATAC-seq of *Foxp1* cKO and control cortices shows disruptions in chromatin accessibility. **A)** Processed snATAC-seq data demonstrates expected cell types and cell distributions per time-point across neocortical development comparable to snRNA-seq data. 9,111 nuclei at E13.5 from N=3/genotype; 5,076 nuclei at E16.5 from N=2/genotype; and 12,620 nuclei at P0.5 from N=3/genotype. **B)** Stacked bar plots show representation from individual replicates across broad cell-type clusters. Star above each bar indicates a significant difference in cell type proportion between genotypes using the two-proportions z-test with Yates correction; green for control and purple for cKO led difference. **C)** Number of differentially accessible regions (DARs) per broad cell-type timepoint highlights shifting role of *Foxp1* regulation. DARs were defined as significant with adj. p-val < 0.05 and avg |log2FC| > 0.25. **D**) Genomic location distribution of each significant DAR peak stratified by closed or open status and time point.

These results indicate a shift in chromatin accessibility due to *Foxp1* loss across the surveyed developmental timepoints. Changes in chromatin accessibility at the regulatory regions of a gene can influence the transcriptional activity of that gene. Overlapping DARs and DEGs indicate a possible regulatory relationship by which changes in chromatin accessibility may contribute to the corresponding changes in gene expression. For example, *Marcks* (myristoylated alanine-rich C-kinase substrate), an actin-cross-linking protein ^39^, is linked to an open DAR in cKOs at E13.5 and an upregulated DEG (Supplemental Tables 1, 2). Moreover, FOXP1 may regulate other transcription factors that recruit chromatin remodeling complexes or histone-modifying enzymes, resulting in changes to chromatin structure and accessibility ^40-42^.

### DEGs confirmed in Foxp1 cKOs are related to neurodevelopmental disorders / ASD

We next sought to validate select DEGs using single molecule fluorescent *in situ* hybridization (smFISH) to maintain cell type specificity. As FOXP1 is relevant to ASD, we elected to validate DEGs with ASD-relevance. Based on our snRNA-seq, we selected three genes: *Rora* and *Nrxn3* are downregulated whereas *Kirrel3* is upregulated in post-mitotic neurons of the upper and deep layers in *Foxp1* cKO mice at P0.5 (Fig. 3A). *Rora* (retinoic acid-related orphan receptor-alpha) is a nuclear steroid hormone receptor and transcriptional activator and is related to the direct regulation of multiple genes associated with ASD ^43^. RORA protein levels are reduced in the postmortem prefrontal cortex and cerebellum of individuals with ASD ^44^. Further, *RORA* is differentially methylated in ASD, leading to its methylation-specific silencing ^44^*. Nrxn3* (neurexin 3) encodes a presynaptic adhesion protein that plays critical roles in synapse formation and function. *NRXN3* variants have been identified in individuals diagnosed with ASD ^45-47^. Some *NRXN3* variants impact synaptic functions and thereby neurodevelopment ^47^. *Kirrel3* encodes a putative cell-adhesion molecule. Variants in *KIRREL3* have been identified in ASD probands and in individuals with ID / NDDs ^48-51^. *Kirrel3* KO mice with ASD-/ID-relevant missense variants show attenuated synaptogenic function ^52^. Further, mice lacking *Kirrel3* are hyperactive ^53^, a phenotype we previously reported in adult *Foxp1* ^+/-^ cKO mice ^29^. Since *Kirrel3* levels are increased, rather than decreased, in *Foxp1* cKOs, this may indicate that overall *Kirrel3* levels are tightly regulated within a specific range for proper function. We independently confirmed these results using smFISH (Fig. 3). We quantified the expression of these genes using their respective RNAscope probes in L5 neurons, where *Ctip2* (*Bcl11b*) was used to mark L5 (Fig. 3B-D), analyzed the total number of pixels per nucleus using a linear mixed model, and confirmed the snRNA-seq findings (Fig. 3E-G). Together, confirmation of these FOXP1-affected genes demonstrates broader impacts on cellular features related to NDDs / ASD and highlights how FOXP1 regulates a network of ASD-relevant genes that impact disease susceptibility and synaptic functions.

**Figure 3:**
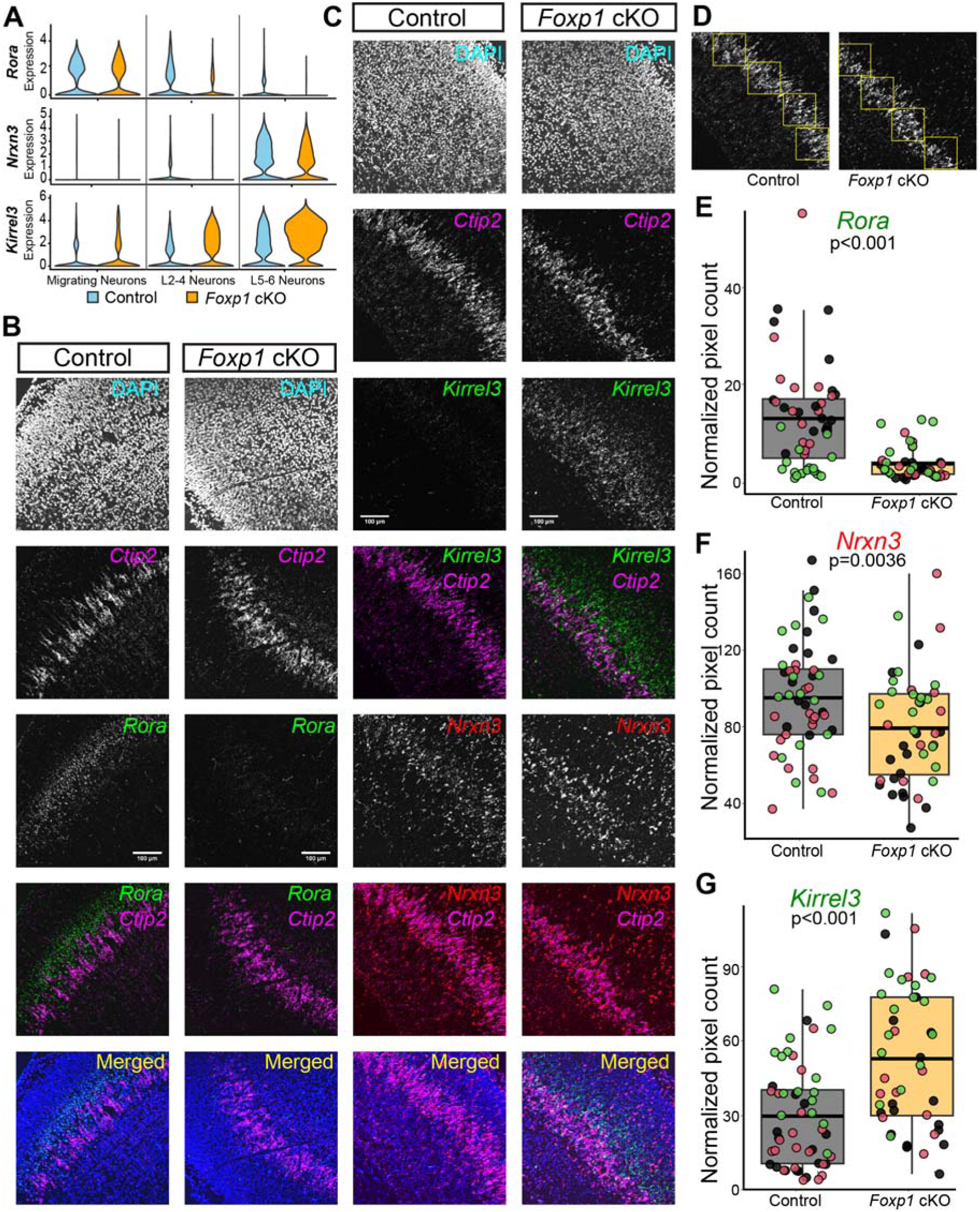
Validation of DEGs identified at P0.5 using RNAScope. **A)** Violin plot demonstrating the expression levels of three representative DEGs by cell type at P0.5. *Rora* and *Nrxn3* are downregulated whereas *Kirrel3* is upregulated. **B, C)** Representative RNA in situ hybridization (ISH) images for *Rora* **(B)** and *Kirrel 3* and *Nrxn3* **(C)** confirm snRNA-seq findings in control and *Foxp1* cKO mice at P0.5. *Ctip2* was used to label L5-6 cortical neurons. **D)** The total number of pixels in each of the 4 bins were measured in the *Ctip2* expressing layer and were normalized to the total nuclei in their respective bins. **E-G)** Box plots illustrate significant downregulation of *Rora* **(E)** and *Nrxn3* **(F)** and upregulation of *Kirrel3* **(G)** in the L5-6 cortical layers of *Foxp1* cKO mice compared to controls. Each dot in the box plots represents normalized pixel counts from individual bins, with colors indicating data from different mice in each group. Significance tested using a linear mixed model with genotype as the fixed factor and individual as the random factor, nested with sections, hemisphere and bins. N = 3 mice per genotype, with 2-3 sections from each mouse. Scale bar:100 µm.

### DEGs and DARs are broadly enriched with ASD-relevant and synaptic genes

To gain further insights into the potential disease-relevant roles of the identified DEGs and DARs, we employed hypergeometric enrichment to assess overlap with ASD datasets (Fig. 4A,B). Specifically, we used SFARI high-confidence ASD-associated genes ^54^, or the top ASD-associated genes identified in two large exome sequencing studies of ASD ^55,56^. Overall, we found that DEGs and DARs across these three developmental timepoints showed enrichment with high-confidence ASD genes.

**Figure 4:**
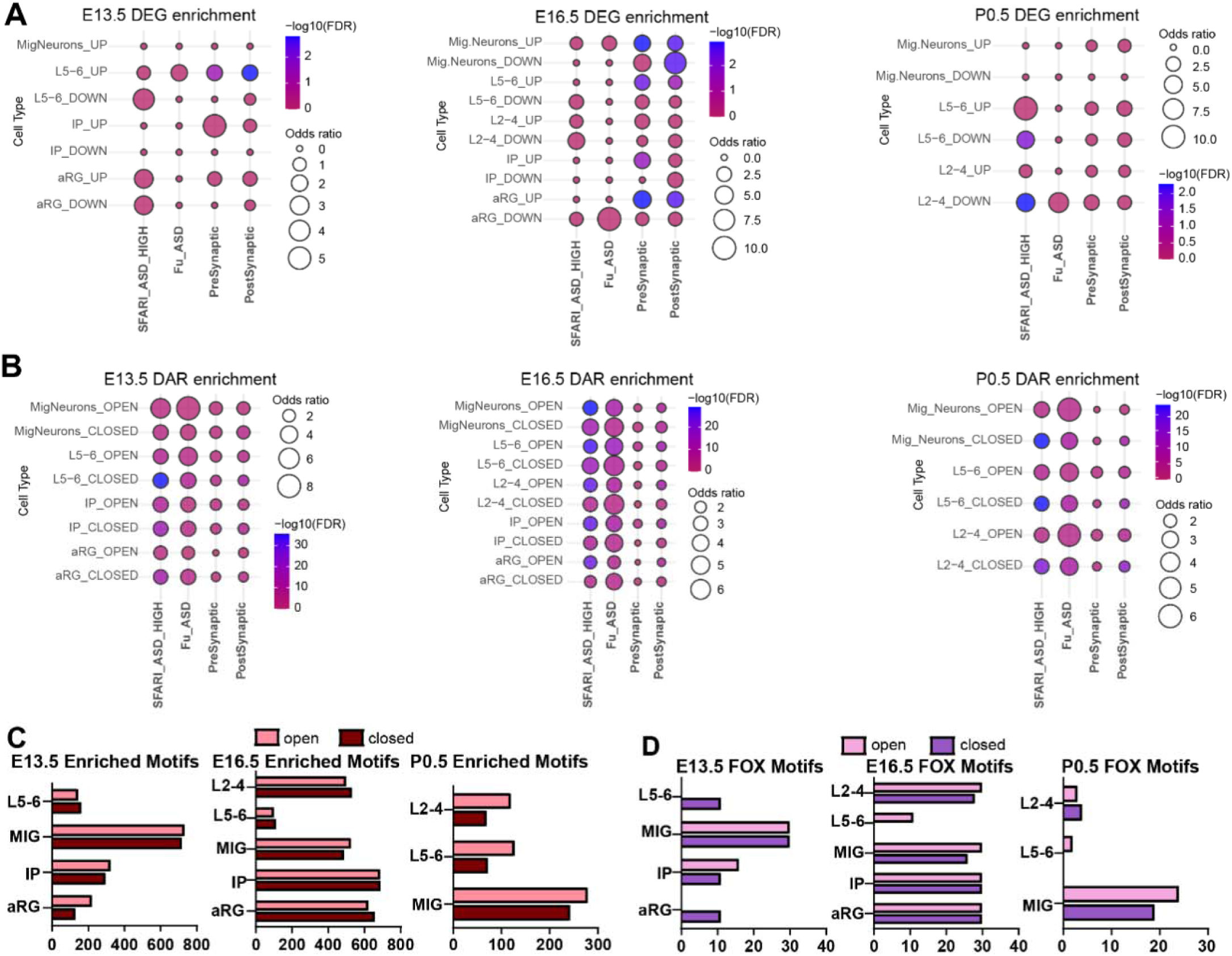
DEGs and DARs are associated with ASD-relevant and synaptic genes and overlapping DEGs/DARs are enriched for FOX motifs. **A, B)** Hypergeometric enrichment FDR and odds ratio output of **(A)** DEGs or **(B)** DARs with high-confidence SFARI-ASD lists, ASD genes from Fu et al., and pre- or post-synaptic genes from Synaptome.db. **A)** In synaptic gene datasets, DEGs have a higher enrichment with post-synaptic genes. **B)** Enrichment is highest for closed DARs and SFARI-ASD genes and synaptic genes at E13.5 and P0.5, with the reverse at E16.5. Hypergeometric enrichment was significant with adj. p-val < 0.05 and - log_10_(FDR)>1.31. Size of circle indicates odds ratio estimated from Fisher’s test. **C)** Number of enriched motifs in open or closed differentially accessible peaks. Defined as significant with adj. p-val < 0.005 and avg |log2FC| > 0.25. **D)** Number of enriched FOX motifs.

Neurons showed a robust association with ASD-relevant genes in terms of both DEGs and DARs. Though we anticipated enrichment between neuronal DEGs and ASD genes based on previous work in both postnatal mice and human brain organoids ^25,57^, we tested whether neural progenitor cells (NPCs) would also share this enrichment since this has not been previously investigated in *Foxp1* cKOs. Both aRGs and IPs also exhibited significant enrichment with ASD genes (Fig. 4A,B), albeit to a lesser extent than neurons. The altered gene expression in NPCs with FOXP1 loss could also contribute to an increased susceptibility to neurodevelopmental disorders (NDDs) since timing of neurogenesis of deep-layer neurons is altered due to variants in select ASD-relevant genes ^58^.

We next sought to investigate DEG and DAR enrichment with synaptically-associated genes ^59^. This was motivated by other work highlighting the role of ASD genes in synaptic function and neuronal activity ^60-63^. Both DEGs and DARs displayed significant enrichment with presynaptic- and post-synaptic-associated genes (Fig. 4A,B; Supplemental Table 3). Interestingly, even DEGs and DARs in the NPCs demonstrated enrichment with synaptic genes. Past work has highlighted a key role for bioelectrical properties of progenitors in neural fate determination and neuronal migration ^64^. However, it is possible that these synaptically-associated genes also perform distinct functions in progenitors in addition to their known synaptic functions. Overall, these findings help underscore a strong association of FOXP1 with synaptic genes. This is important to highlight given that synaptic transmission is commonly disrupted in other ASD models ^62^.

### Potential direct FOXP1 targets revealed by FOX motifs, integration with FOXP1 ChIP-seq, joint snRNA-seq/snATAC-seq analysis

To assess the functional significance of DARs, we searched for enriched transcription factor (TF) motifs to infer potential cell type and age-specific transcriptional regulation (Fig. 4C, Supplemental Table 2). We identified enriched TF motifs across all timepoints in both open and closed DARs. Notably, migrating neurons exhibited the highest number of enriched motifs, while deep layer (DL) neurons showed the fewest (Fig. 4C). Additionally, the embryonic timepoints demonstrated more motif enrichment than the postnatal timepoint (Supplemental Table 2). We then specifically investigated the presence of enriched FOX family motifs (Fig. 3D). Enriched FOX motifs were observed at all timepoints, except at E13.5 in open aRG cells or L5-6 neurons, and at E16.5 and P0.5 in closed L5-6 neurons (Fig. 4D, Supplemental Table 2).

Recent work used chromatin immunoprecipitation sequencing (ChIP-seq) to identify the regulatory targets of select ASD-associated genes, FOXP1 included, in the mouse cortex ^65^. Using our snATAC-seq data, we found high overlap between the peaks in that study and our snATAC-seq DARs (Supplemental Fig. 5A-E). We specifically overlapped our E16.5 snATAC-seq DARs with E15.5 ChIP-seq peaks and P0 snATAC-seq DARs with E18.5 ChIP-seq peaks (Supplemental Fig. 5A). We show feature distribution for the shared features (Supplemental Fig. 5B-C). Further, GO terms for overlapping E16.5 snATAC-seq DARs with E15.5 ChIP-seq peaks are related to axonogenesis, neuron differentiation, axon guidance, and synapse assembly (Supplemental Fig. 5D). For P0 snATAC-seq closed DARs overlapping with E18.5 ChIP-seq peaks in L5-6 neurons, GO terms include axonogenesis, neurogenesis, ubiquitination, and corpus callosum development (Supplemental Fig. 5E, Supplemental Table 3). Interesting, a thinner corpus callosum or reduced corpus callosum volume is a phenotype previously observed in juvenile and adult *Foxp1* cKO mice ^29,30^. Additionally, the corpus callosum is often impacted in individuals with ASD ^21,66^.

Next, we sought to identify genes that vary between genotypes driven by a joint snRNA-seq and snATAC-seq analysis. To do so, we used GLUE (Graph-Linked Unified Embedding) ^67^ to match snRNA-seq and snATAC-seq cells, allowing for integration of our single-cell multi-omics data. Then we applied MOFA+ (Multi-Omics Factor Analysis v2) ^68^ and decoupler to do a factor analysis to find features that vary between conditions, i.e., genotypes. Each factor captures the coordinated gene expression across cell types ^68^. This approach identified significant factors (which capture most of the variance observed) and top loadings, or genes that are most important for driving each factor. We then performed GO enrichment analyses for top genes in statistically significant factors (Supplemental Fig. 5F-I). At E13.5, GO terms for aRGs included neuron differentiation, pallium development, and morphogenesis involved in differentiation (Supplemental Fig. 5F). At E16.5, GO terms for L2-4 neurons include cell adhesion via plasma membrane adhesion molecules, tubulin binding, neurogenesis, and cell cycle transition. Some top driver genes include *Robo1*, *Tubb3*, and *Kirrel3* (Supplemental Fig. 5G). At P0, enriched GO terms in L5-6 neurons and L2-4 neurons relate to axonogenesis, plasma membrane adhesion, neuron development, and differentiation (Supplemental Fig. 5H-I). In L5-6 neurons at P0, top genes that vary between genotypes in the joint snRNA-seq and snATAC-seq analysis include *Robo1*, *Nrxn3*, and *Cdh10* (Supplemental Fig. 5H). Genes that vary between genotypes in L2-4 neurons at P0 include *Kirrel3*, which is a DEG we confirmed (Fig. 3), *Cux1*, *Epha3*, and *Sema3a* (Supplemental Fig. 5I).

### Loss of Foxp1 results in a more mature pseudotime developmental trajectory at E13.5

We next specifically focused on the role of FOXP1 at E13.5, because at this timepoint FOXP1 is expressed in both NPCs and post-mitotic neurons (Supplemental Fig. 3). We sought to determine how FOXP1 differentially regulates the developmental trajectory of each of these cellular populations at E13.5. We used the *CellType_coll* annotation which consists of broad cluster per cell type for ease of visualization (Fig. 5A). We then performed pseudotime analysis using diffusion pseudotime (see Methods)^69^. As pseudotime infers an ordering for each cell along a lineage based on gene expression, from immature to mature, we can compare changes due to *Foxp1* deletion along a differentiation trajectory. This allows us to reconstruct dynamic gene expression programs underlying biological processes.

**Figure 5:**
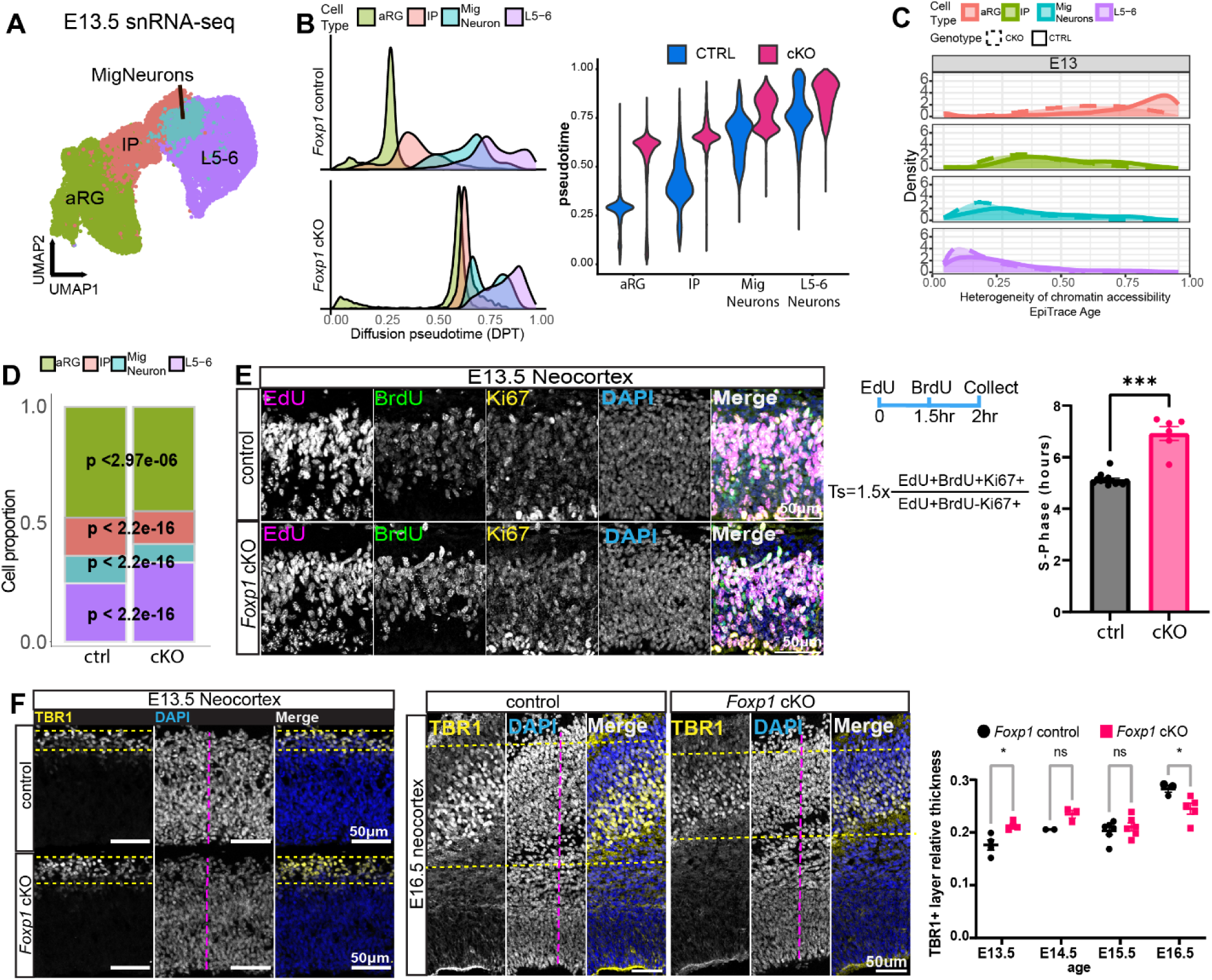
*Foxp1* cKO results in neurons with accelerated pseudo-age and precocious neurogenesis. A) UMAP plot of E13.5 snRNA-seq data with excitatory neuronal lineage broad cell-type annotation. **B)** *(Left)* Pseudotime plot of cell-type density distribution vs pseudotime. Path lengths between control are significantly different p < 2e-16 with Mann-Whitey test with continuity correction. *(Right)* Violin plot showing pseudo-age by genotype and cell types. **C)** EpiTrace estimate of cell replicational age indicates aRGs in *Foxp1* cKOs have a younger cell replication age than controls. **D)** Stacked bar plot of cell-type proportion by genotype of E13.5 snRNA-seq data. p-values of two-proportions z-test with Yates correction. *Foxp1* cKOs have significantly reduced proportion of aRGs, IPs, migrating neurons, and significantly higher proportion of L5-6 neurons. N=3/genotype. **E)** Estimation of S-Phase length shows an elongated S-Phase length in *Foxp1* cKOs at E13.5. N=9/ctrl and 6/cKO. Mann-Whitney test. **F)** IHC images at 20x of TBR1+ layer thickness relative to DAPI+ thickness at E13.5 (*left*) and E16.5 (*middle*) and quantification of TBR1+ relative layer thickness from E13.5-E16.5 (*right*). TBR1 layer thickness at E13.5 is greater in cKOs at E13.5 but reduced relative to controls by E16.5. Replicates by age are E13.5: N=4; E14.5: N=2-3; E15.5: N=6; E16.5: N=3-5 per genotype. Mean±SEM. Scale bars = 50μm. apical radial glia (aRG); intermediate progenitors (IPs); Mig Neurons (Migrating neurons); L5-6 (L5-6 neurons).

We report that the pseudotime distribution of cell types in *Foxp1* cKOs is condensed compared to the control mice, with reduced separation between cell transitions, especially between aRGs and IPs (Fig. 5B, Supplemental Fig. 6A). Overall, the pseudo-age value for all cKO cell types is older than that assigned to control cells as shown in terms of distribution (Fig. 5B *left*, Supplemental Fig. 6A). We also show diffusion pseudotime of the cell types in a reduced dimensional space DC1xDC2, calculated in Destiny, colored by cell type and pseudo-age (Supplemental Fig. 6A). Further, the total path length between the cKO and control developmental trajectories is significantly different (Mann-Whitney test with continuity correction adj. p-val < 2.2e-16). Therefore, FOXP1 loss leads to an estimated older cellular pseudo-age, indicating that FOXP1 plays a role in regulating the development of cells during early neurogenesis.

### Loss of Foxp1 alters gene expression to reduce progenitor maintenance and promote differentiation

To determine the signaling pathways downstream of FOXP1 driving changes in the developmental trajectories of cell types, we assessed the functional enrichment of gene ontology (GO) categories within the DEGs/DARs of aRGs and DL neurons (Supplemental Fig. 6C,D; Supplemental Table 4). Upregulated DEGs in DL neurons at E13.5 are enriched for GO terms for regulation of neuron differentiation, DNA replication, negative regulation of Notch signaling pathway, and telencephalon development (Supplemental Fig. 6C, Supplemental Table 4). GO terms for open DARs in aRGs represent positive regulation of cell differentiation, chromatin remodeling, Wnt signaling, and regulation of microtubule (de)polymerization (Supplemental Fig. 6D *left*, Supplemental Table 4). Closed DAR GO terms involved regulation of cell migration, transcription co-repressor activity, ephrin receptor activity, and transmembrane receptor protein tyrosine kinase activity (Supplemental Fig. 6D, *right*; Supplemental Table 4). These terms highlight changes in the biological function of aRGs and the underlying mechanisms driving observed changes in developmental trajectories. These results suggest that FOXP1 represses genes associated with aRG differentiation to permit self-proliferation and therefore, at E13.5, aRGs in cKOs express more pro-differentiation genes, resulting in an advanced estimated pseudo-age.

snRNA-seq data can reveal changes not only in differential expression but also in cell type proportions. At E13.5, we observed significant shifts in cell type proportions in cKOs, with significantly fewer aRGs (two-proportions z-test with Yates correction ^22^; adj. p-val <2.97e-06) together with a significant relative increase in L5-6 neurons (two-proportions z-test with Yates correction; adj. p-value < 2.2e-16, Fig. 5D) ^22^. Other work reported an increased prevalence of neurogenic-type divisions in radial glia of mice lacking *Foxp1* ^32^. Together, these findings suggest that the loss of FOXP1 promotes neuronal differentiation by altering gene expression and cell type proportions at E13.5.

### Heterogeneity of chromatin accessibility on clock-like loci is reduced in Foxp1 cKOs indicating a younger mitotic / cell replicational age

As an additional measure to determine cell age, we used EpiTrace, a tool that quantifies the fraction of opened “clock-like” loci in snATAC-seq data to estimate the mitotic age or developmental potential of cells ^70^. EpiTrace harnesses previous work showing that age-associated DNA methylation changes occur at specific genomic regions ^70^. As cells divide, the heterogeneity of chromatin accessibility at these clock-like loci is reduced, providing a measure of mitotic age. Applying EpiTrace to the excitatory neuronal lineage at E13.5, we find that aRGs in *Foxp1* cKOs exhibit a younger cell replicational age, indicating fewer mitotic divisions than in controls (Fig. 5C). This observation aligns with our finding of increased pseudo-age/differentiation potential in these cells in *Foxp1* cKOs.

### S-Phase length is increased in progenitors upon loss of FOXP1

To determine the mechanism of improper maintenance of the progenitor pool, we use two distinguishable thymidine analogues, BrdU (5-bromo-2’-deoxyuridine) and EdU (5-Ethynyl-2′-deoxyuridine), to estimate the length of S-Phase at E13.5 ^71^. By co-staining with KI-67, a marker of proliferating cells, we find that NPCs in *Foxp1* cKOs have a significantly lengthened S-Phase (Fig. 5E) and cell cycle length (Supplemental Fig. 6B).

To determine whether other aspects of the cell cycle are impacted by FOXP1 loss, we first used phosphorylated histone M3 (PH3) staining as a proxy of M-Phase. We detected no significant differences in PH3+ cells between cKOs and controls at E13.5 (Supplemental Fig. 6F). DEGs at E13.5 were enriched for GO terms related to protein ubiquitination (Supplemental Fig. 6C). Although apoptotic pathways are known to be tightly regulated by post-translational modifications such as ubiquitination ^72^, we did not detect a change in apoptosis as measured by cleaved-caspase-3 between genotypes at E13.5 (Supplemental Fig. 6G). Thus, the progenitor pool in cKOs is altered collectively through fewer aRGs (Fig. 5C), a longer S-Phase, and dysregulation of genes necessary for progenitor maintenance.

### Precocious generation of DL neurons

The altered developmental trajectory and younger cell replication age of aRGs in cKOs at E13.5 prompted us to further examine their shifting neurogenic potential. Our previous work demonstrated cyto-architectural changes in the P7 cortex of *Foxp1* cKOs, where we observed a relatively thinner cortex overall with reduced DL thickness but a relatively thicker L2-4 ^30^. At E13.5, we found that *Foxp1* cKOs exhibit a relatively thicker TBR1+ cell layer (a marker of DL neurons) compared to controls (Fig. 5F). However, this difference was not observed at E14.5 or E15.5 and by E16.5, the thickness was reduced (Fig. 5F). These results may be attributed to an increased number of differentiative divisions in the NPCs at E13.5 in *Foxp1* cKOs ^32^. Increased neurogenic division may lead to a depletion of the progenitor pool, resulting in an overall reduction in neuron production that is observed postnatally in cKOs.

### Accelerated cell cycle exit from E15.5 to E16.5 with loss of FOXP1

Corticogenesis is characterized by the slowing down of the cell cycle in NPCs and an increased frequency of differentiative divisions, resulting in a higher rate of neurogenesis ^73^. In *Foxp1* cKOs, corticogenesis initially appears to be accelerated (Fig. 5B). To investigate whether the reversal in the thickness of the TBR1+ layer at E16.5 is due to altered neurogenesis, or changes in the cell cycle exit rates at one of the intermediate time-points, we used EdU that incorporates into the cells undergoing DNA replication during S-phase. To measure changes in cell cycle exit, we injected EdU once at E15.5 and analyzed changes after 24 hours. We quantified cells that were cycling at the time of injection (EdU+) but were no longer proliferative 24 hours later (KI67-), indicating they had exited the cell cycle (EdU+Ki67- / EdU cells) (Fig. 6A) ^74^. We observed that cKOs exhibit a higher proportion of cells exiting the cell cycle compared to controls (Fig. 6A). The faster cell cycle exit of NPCs suggests a bias toward neuron production at the expense of their self-renewing divisions. This could be an underlying cause of the relatively thicker L2-4 reported in *Foxp1* cKOs at P7, despite an overall thinner neocortex, as most of the neurons born at E16.5 acquire UL identities ^9,30^. In fact, our snRNA-seq data showed a relative increase in L2-4 neuron proportions in cKOs at P0.5 (Fig. 6B *left*), but not at E16.5 (Fig. 6B *right*) based on the two-proportions z-test with Yates correction ^22^. Further, using EpiTrace ^70^ to infer cell replication age in a cell-type-specific manner at E16.5, we find that L2-4 neurons in cKOs exhibit an older mitotic age (Fig. 6C). Previous studies using EpiTrace have shown that forced differentiation increases the “age” of differentiated cells ^70^. These findings support the idea that neuronal precursors are generating L2-4 neurons faster in cKOs and exhausting their progenitor pool.

**Figure 6:**
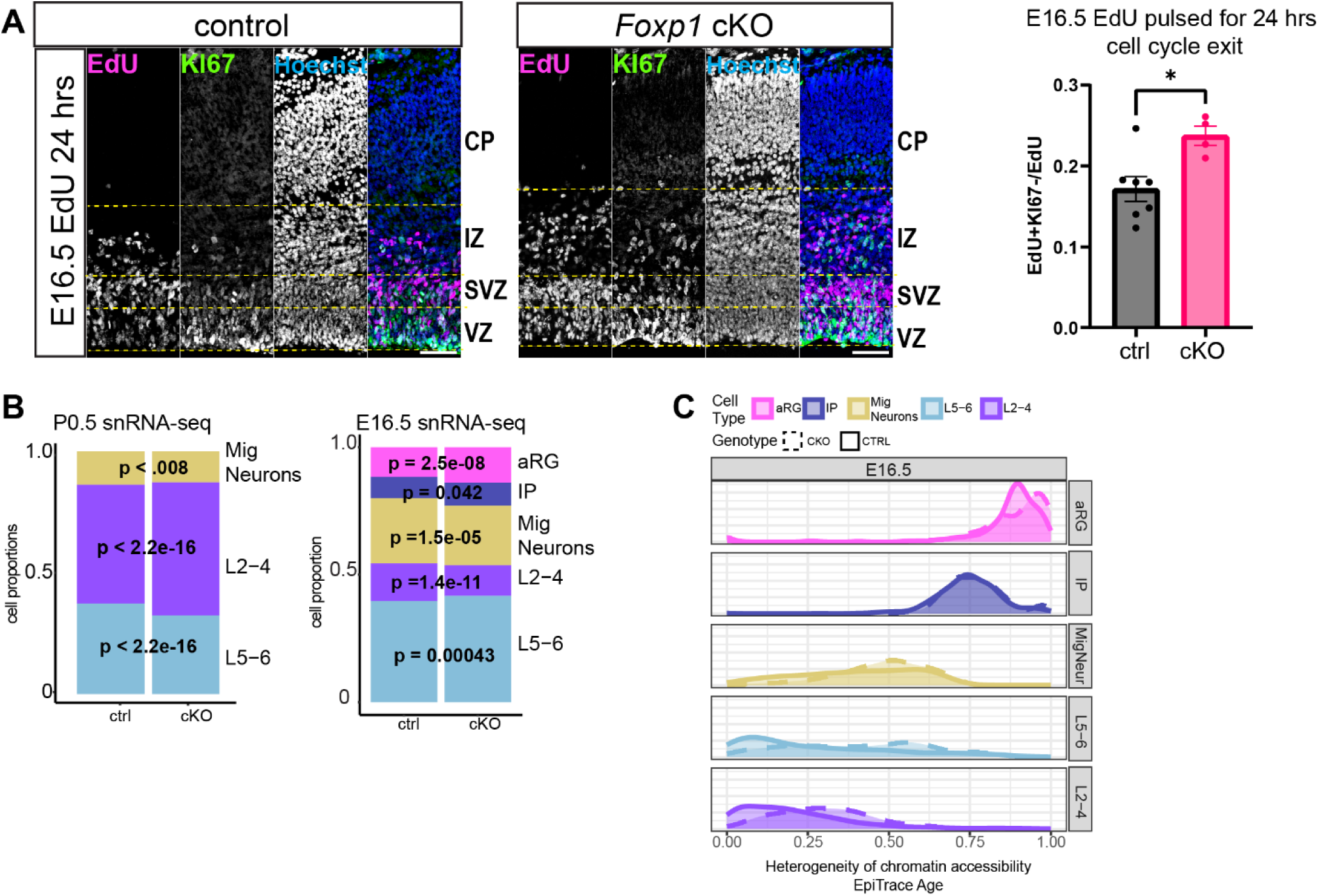
Accelerated cell cycle exit at E16.5 in *Foxp1* cKOs and altered cortical layer marker gene expression. **A)** *(Left*) IHC images show cell cycle exit assay of 24hr EdU pulse from E15.5 to E16.5. KI67: proliferative marker; Hoechst used as DNA marker. *(Right)* EdU+KI67-cells over total EdU cells; Cells that are EdU+ were cycling at the time of EdU injection. KI67-cells are no longer proliferating. EdU+KI67-cells indicate cells that have exited the cell cycle. *Foxp1* cKOs have higher EdU+Ki67-/EdU cells. Mean±SEM. N=4-7/genotype. Mann-Whitney test. VZ: ventricular zone. SVZ: sub-ventricular zone, IZ: intermediate zone, CP: cortical plate. **B)** *left*) Cell-type proportions at P0.5 for snRNA-seq data by broad cell type per genotype. Faster cell-cycle exit during the typical generation of UL neurons (as in panel A) may underlie the postnatal phenotype of relatively thicker L2-4, and relatively thinner L5-6. (*right*) However, these differences are not detected at E16.5 in the snRNA-seq data likely since neuronal identity is still being refined. **C)** EpiTrace age results from E16.5 by cell type. Heterogeneity of chromatin accessibility / division age suggests that L2-4 neurons in cKOs have been generated from precursor cells with greater mitotic age vs controls. Scale bar = 50μm.

### Neuronal migration deficits with loss of FOXP1

Previous work has shown that knock-down of *Foxp1* via *in utero* electroporation results in widespread neuronal migration deficits ^23,28^. However, genetic knock-down of *Foxp1* does not readily reflect such a stark phenotype, but rather a more restricted change, in the postnatal neocortex with mispositioning of CUX1+ cells, a traditional marker of L2-4 neurons, in the DLs at P7 ^30^. Since FOXP1 is also present in the NPCs, loss of FOXP1 may affect neuronal fate. Therefore, we investigated if the ectopic CUX1+ cells were a result of migration deficits or due to changes in cell fate specification. We pulsed EdU at 15.5 and analyzed brains at P7 and P16.5. We co-stained the brain sections with CUX1 (L2-4 marker) and CTIP2 (L5-6 marker) to identify EdU+CUX1+ cells in the DL (Fig. 7A). We found EdU+CUX1+ cells in the DLs at P7 (Fig. 7A,B), but we did not observe this phenotype if EdU was injected at earlier time-points (data not shown). The majority of neurons generated at E15.5 are destined to form UL neurons ^9^. We quantified the number of EdU+ cells in the UL and DL and determined that *Foxp1* cKOs have a significantly higher number of EdU+ cells in the DLs compared to controls (Fig. 7C), suggesting a migration deficit phenotype. To investigate if there was a delay in migration, we assessed these effects at P16.5. We still observed EdU+/CTIP2-cells in the DL (Fig. 7D,E; Supplemental Figure 7A-B). As the number of EdU+ cells in the DLs remains significantly higher in cKOs, this indicates a primary migration deficit, not a delay in neuronal migration (Fig. 7E). Further, DEGs and DARs at E16.5 and P0.5 also highlight genes with key functions in neuronal migration, further supporting dysregulation in migration cues (Supplemental Fig. 7C, Supplemental Table 4). Overall, our results suggest that UL neurons are being generated at the anticipated times but are unable to migrate to their expected destination in the *Foxp1* cKOs.

**Figure 7:**
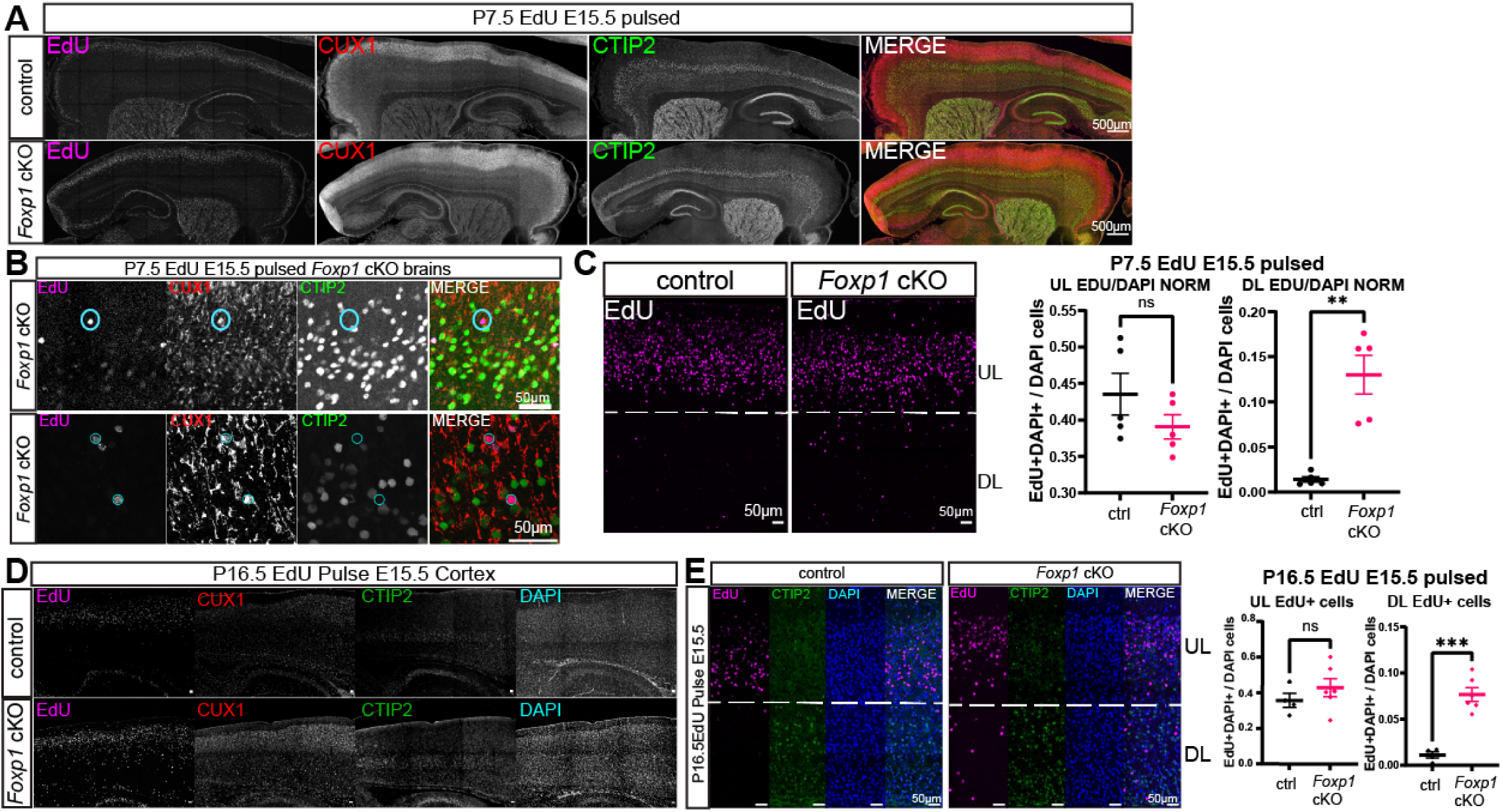
Selective neuronal lineage migration deficits but no change in timing of neurogenesis in *Foxp1* cKO animals. **A-B)** EdU birth-dating experiment shows that neurons generated from progenitors dividing at E15.5 remain in the DLs in *Foxp1* cKOs at higher levels than in controls throughout the cortex. Some of these cells are EdU+CUX1+ indicating they have an UL cell type identity and are CTIP2-. **B)** *Foxp1* cKO cortices *(top)* is at 20x and (*bottom*) at 63x showing EdU+CUX1+CTIP2-cells in L5-6 region. **C)** P7.5 brains stained for EdU that was pulsed at E15.5 have significantly more cells in the DLs in *Foxp1* cKOs. No significant difference between EdU+ cells in the ULs between genotypes. N=5/genotype. **D)** P16.5 brains from E15.5 EdU pulsed animals with CUX1 and CTIP2 to demarcate cortical layer boundaries at 10x resolution. **E)** P16.5 brains with EdU cell quantification in ULs or DLs shows a significant number of EdU+ cells located in the DLs in cKOs but not controls. No significant change in the number of UL EdU+ cells. N=5/genotype. EdU+ cells were CTIP2-despite being in L5-6, suggesting appropriate timing of neurogenesis at E15.5 for UL cells and a likely selective migration deficit that remains at P16.5. Mann-Whitney test. Mean±SEM shown.

## Discussion

Our study revealed cell type-specific changes in gene expression and chromatin accessibility changes during early cortical development in mice with *Foxp1* loss. FOXP1 is expressed in both NPCs and post-mitotic neurons at E13.5. Upon FOXP1 loss in NPCs, we observed faster generation of DL neurons. However, this precocious development of DL neurons depleted the progenitor pool, leading to a reduction in the relative proportions of DL neurons. We also found a faster cell cycle exit in NPCs by E16.5. Additionally, migration and expression of canonical layer-specific TFs were disrupted in DL neurons. Finally, these cellular phenotypes occur in parallel with molecular changes in gene expression relevant to synaptic function and NDD pathophysiology.

### Selective migration deficits of a subtype of upper layer neuron

Previous research has highlighted the importance of FOXP1 in regulating neuronal migration ^23,28,30^. Our method of birth-dating neurons in *Foxp1* cKO mice using a thymidine-analogue allowed us to definitively disentangle a migration impairment from a cell fate specification phenotype. We opted to use a thymidine analogue pulse, although other techniques exist, such as FlashTag ^75^, which tag dividing progenitors that contact the ventricular surface to mark a more precise group of cells. Thymidine analogues incorporate their mark into DNA of all progenitors and their progeny during DNA replication in S-Phase. Though this method labels a broader group of cells, we could observe patterns of cell generation based on date of injection and track generation of neurons and their ultimate position based on birthdate.

Using single cell genomics, we also obtained unbiased genomic profiling of cells at each timepoint. As an example, at P0.5, *Rorb* (*RAR-related orphan receptor B*) is a downregulated DEG and closed DAR in L2-4 neurons (Supplemental Table 1). Previous research reported that individuals with *RORB* mutations exhibit mild-to-moderate ID and seizures ^76^. Since almost 12% of individuals with FOXP1 syndrome also exhibit seizures ^21^, this suggests a possible connection between RORB and FOXP1 with respect to seizure susceptibility. Furthermore, changes in *Rorb* expression were observed in a cell population that neither expresses FOXP2 nor shows FOXP4 upregulation (Supplemental Table 1). This suggests these cells are more vulnerable to the loss of FOXP1, given the lack of compensation by these paralogous genes.

### FOXP1 regulates genes relevant for ASD

*FOXP1* is a high confidence ASD-associated gene and previous studies have shown that FOXP1 regulates genes important for neurodevelopment ^25,30,55,56^. We specifically highlight increased enrichment of DEGs and DARs with ASD genes across all timepoints. Given the heterogeneity of genetic mutations and molecular mechanisms contributing to ASD, it is important to understand convergent mechanisms for ASD-associated genes. For example, previous research seeking to coalesce changes due to ASD risk genes found asynchronous maturation of DL neurons and interneurons ^58^. Here, we report changes in molecular function and generation of DL neurons and additional changes in UL neurons with the loss of FOXP1 (Fig. 7B). Thus, these data may be helpful to gain additional insight into the shared changes in UL neurons in other models assessing ASD risk genes.

Further, we observed distinct patterns between the number of open/closed DARs and upregulated/downregulated DEGs from E16.5 to P0.5 (Fig. 1C, 2C). As FOXP1 itself is not known to be a chromatin modulator, these data may suggest it is involved in the regulation of a network of genes that are chromatin modulators. This may result in more accessible regions that encode transcriptional repressors, affecting the number of upregulated DEGs in cKOs. As development is dynamic, changes in patterns of accessibility and gene expression may vary across ages. Thus, it may be beneficial to study additional ages with added modalities (e.g. ChIP-seq) to better understand how FOXP1 impacts gene regulation.

A recent study used ChIP-seq to identify the regulatory targets of five ASD-associated transcriptional regulators, including FOXP1 ^65^. The authors determined that these regulators share substantial overlaps in binding sites within other ASD-associated genes ^65^. We find a substantial overlap of our snATAC-seq DARs and their data. The reduction of either ARID1B or TBR1 expression (two regulators included in the ChIP-seq study) in cultured neonatal mouse cortical cells resulted in a reduction of neuronal (NeuN+) cells, similar to the phenotype we observed in postnatal *Foxp1* cKO brains ^65^. Together, these data underscore the importance of conditional knock-out studies to better understand the role of cortical FOXP1 in convergent neurodevelopmental disorder outcomes.

### FOXP1 regulates genes encoding synaptic proteins across development

We also determined that both the DEGs and DARs in cKOs are enriched for synaptic protein encoding genes, especially in postnatal tissue. Though changes in the expression of genes encoding synaptic proteins are present from early development, their persistence and increasing enrichment among DEGs over development validates their postnatal relevance. This may also implicate changes in the micro- and macro-circuity of the developing neocortex. Our recent work has highlighted the importance of FOXP1 in regulating potassium currents in striatal neurons ^77^, which underscores the possibility of a similar role in excitatory cortical neurons.

## Conclusions

Our data provide insights into the *in vivo* role of FOXP1 across early neocortical development at cellular resolution. We show that FOXP1 is critical for the function of NPCs, impacting the rate of neurogenesis indirectly by altering their proliferation. We also demonstrate how *Foxp1* deletion directly impacts cortical migration. Moreover, we identify the DEGs and DARs in *Foxp1* cKO mice that are enriched for genes encoding ASD and synaptic proteins and may underlie the cell type-specific phenotypes. We also report complementary outcomes while analyzing the snRNA-seq and snATAC-seq individually in terms of cellular maturation. These findings provide opportunities for cellular and molecular manipulations to rescue relevant phenotypes associated with FOXP1 syndrome and other forms of ASD.

## Limitations of the study

The data in this study present molecular insights into the *in vivo* roles of FOXP1 across select early timepoints during early neocortical development. These timepoints were selected based on the expression patterns of FOXP1 and postnatal ASD-relevant phenotypes. However, it is important to recognize that neurodevelopment occurs rapidly in the murine brain. Thus, this study provides only select snapshots during early neurogenesis, late neurogenesis, and when FOXP1 has layer-restricted expression at one postnatal timepoint. Further, we lose spatial information using snRNA-seq and thus cannot identify the altered gene expression in the select neurons that exhibit migration deficits. Also, this work focuses on the effects of conditional *Foxp1* deletion in the excitatory neuronal lineage driven by the *Emx1.*Cre, which was motivated by our previous work. However, FOXP1 is expressed in additional cell types, which may also exhibit non-cell-autonomous effects due to *Emx1*.Cre driven *Foxp1* deletion. Thus, this work explores the effects of FOXP1 loss in cells outside of the excitatory neuronal lineage to a limited capacity. However, additional investigations into such cell types would be informative. Future studies may also benefit from exploring additional time points to gain a more comprehensive understanding of how FOXP1 regulates neocortical development, especially using a spatially preserving approach that provides cellular resolution. This is particularly salient as another study has found that FOXP1 affects transcriptional programs regulating angiogenesis and glycolysis during early neurogenesis at E12.5 ^78^. Finally, while we uncover ASD-relevant gene expression programs downstream of FOXP1 using *Emx1^Cre/+^;Foxp1^flox/flox^* conditional deletion, a FOXP1-syndrome-relevant haploinsufficient mouse model may reveal additional or different downstream pathways.

## Resource availability

### Lead contact

Further information and requests should be directed to and will be fulfilled by the Lead Contact, Genevieve Konopka.

### Materials availability

Animals and materials generated from this study are available from the lead contact with a completed Materials Transfer Agreement.

### Data and Code Availability

- The sequencing data reported in this paper can be accessed at NCBI GEO with accession number: GSE267673 (GSE267670, GSE267671).

(https://www.ncbi.nlm.nih.gov/geo/query/acc.cgi?acc=GSE267673) All other acquired data are available upon request to the Lead Contact.

- Code that was used to perform data pre-processing, clustering, differential gene expression analysis, and differential accessibility region analysis is available at the GitHub repository: (https://github.com/anabrains/neocortex_genomics_Foxp1)
- Any additional information required to reanalyze the data reported in this paper is available from the Lead Contact upon request.

## Acknowledgments

Our sincerest thanks to Rachael Vollmer and Drs. Yuxiang Liu and Willy R. Vasquez for providing feedback on the manuscript. G.K. is a Jon Heighten Scholar in Autism Research and Townsend Distinguished Chair in Research on Autism Spectrum Disorders at UT Southwestern. This study was funded by UTSW T32 Molecular Medicine Research Training Program (NIH T32GM109776), R01MH102603 diversity supplement, F31MH123140-01, and HHMI Gilliam Fellowship to A.O; F30DC022213 to M.J.; the NIH (R01MH102603, R01MH126481), and the Simons Foundation for Autism Research Award (573689) to G.K. We also thank the Neuroscience Microscopy core at UT Southwestern.

## Author Contributions

A.O. and G.K. designed the study. G.K. edited the manuscript. A.O. and F.A. collected and processed tissue for snRNA-seq. M.H. maintained mouse lines and performed genotyping. NK collected tissue for and performed the RNAscope RNA *in situ* hybridization and quantification. EO performed the EpiTrace analysis and the GLUE/MOFA+ analysis. M.J. assisted with mouse husbandry and immunohistochemistry. A.O. performed mouse husbandry; collected, and processed tissue for immuno-histochemistry, snATAC-seq, performed bioinformatic analyses, and wrote the manuscript.

## STAR⍰Methods

### Key Resources table

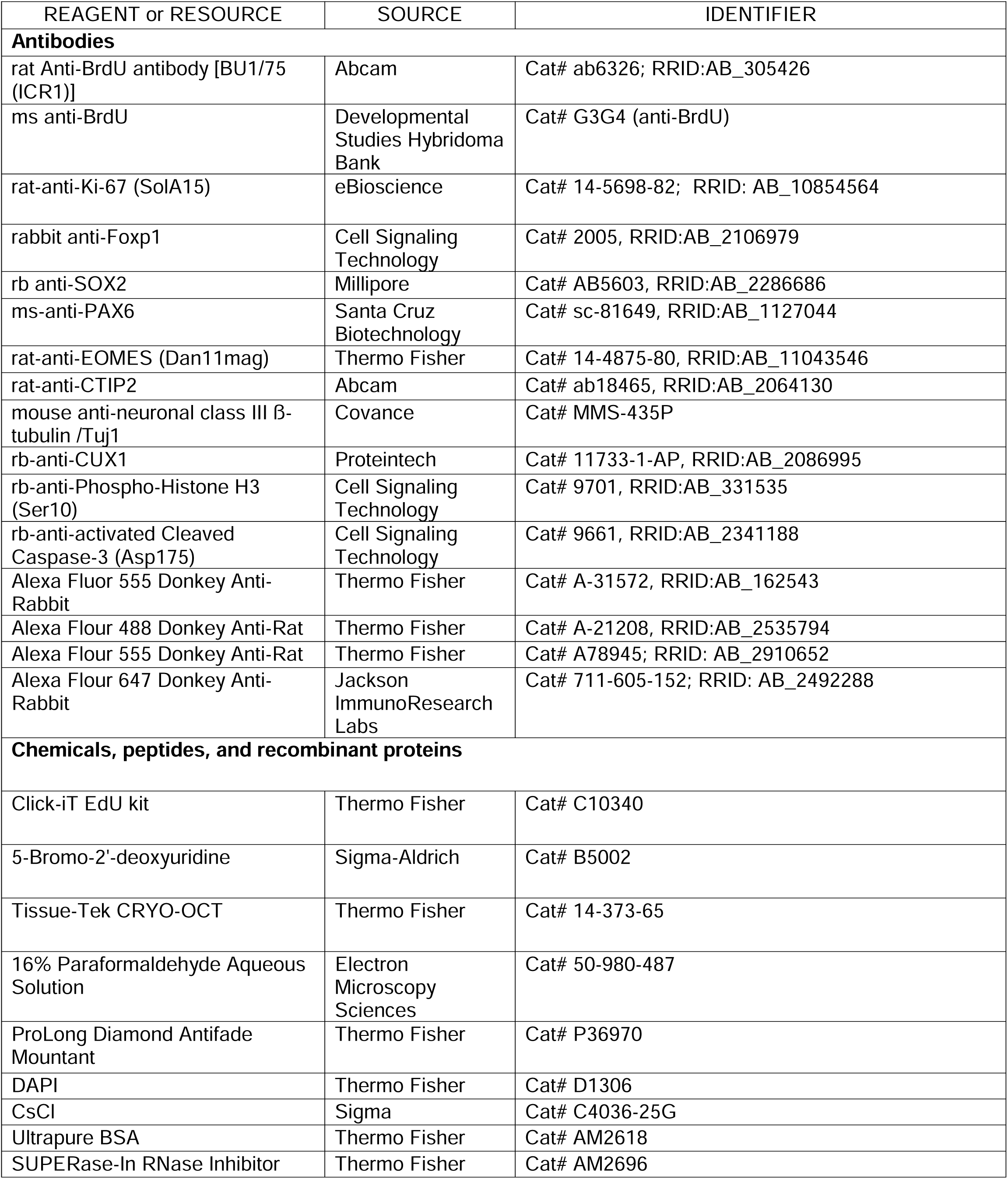
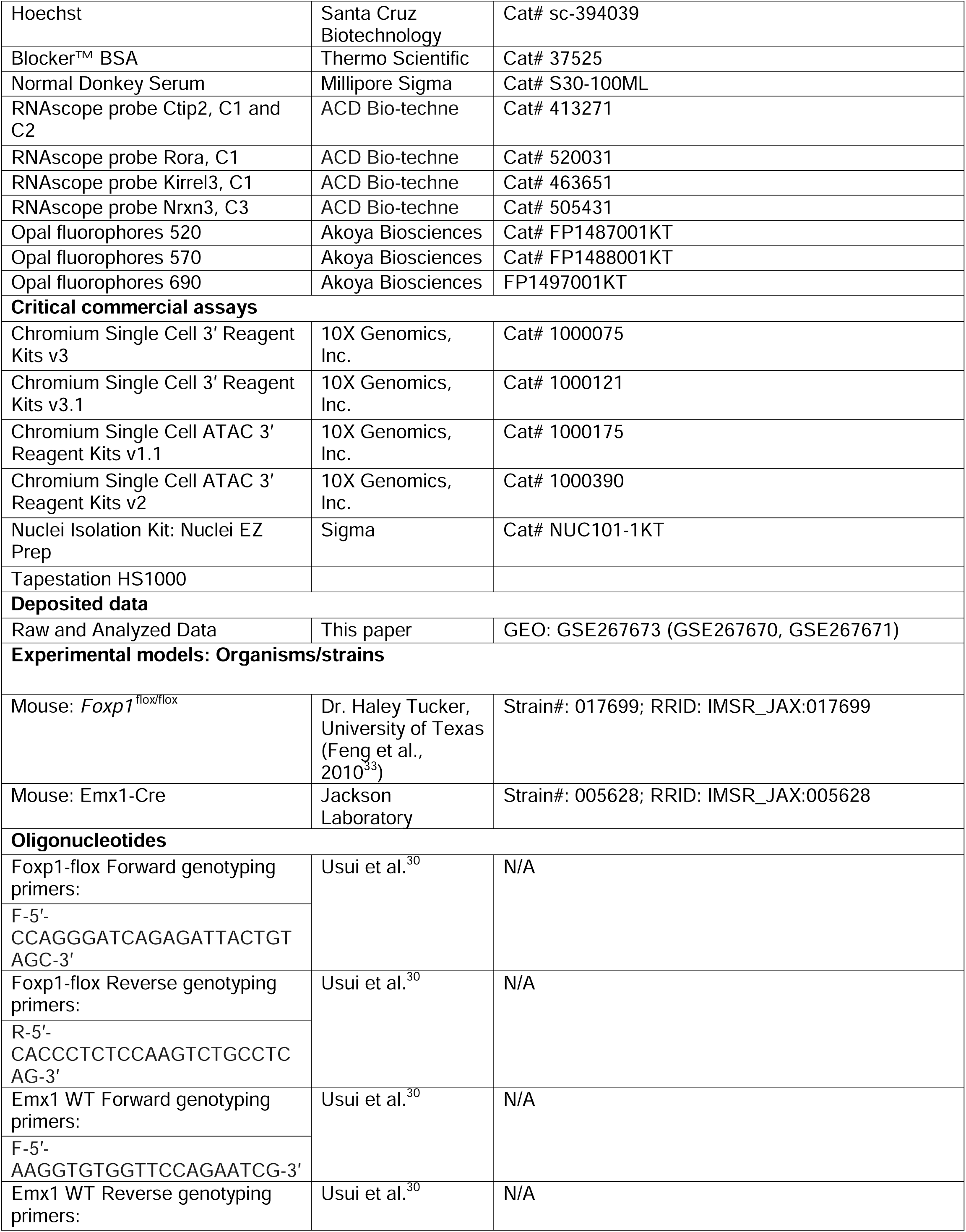
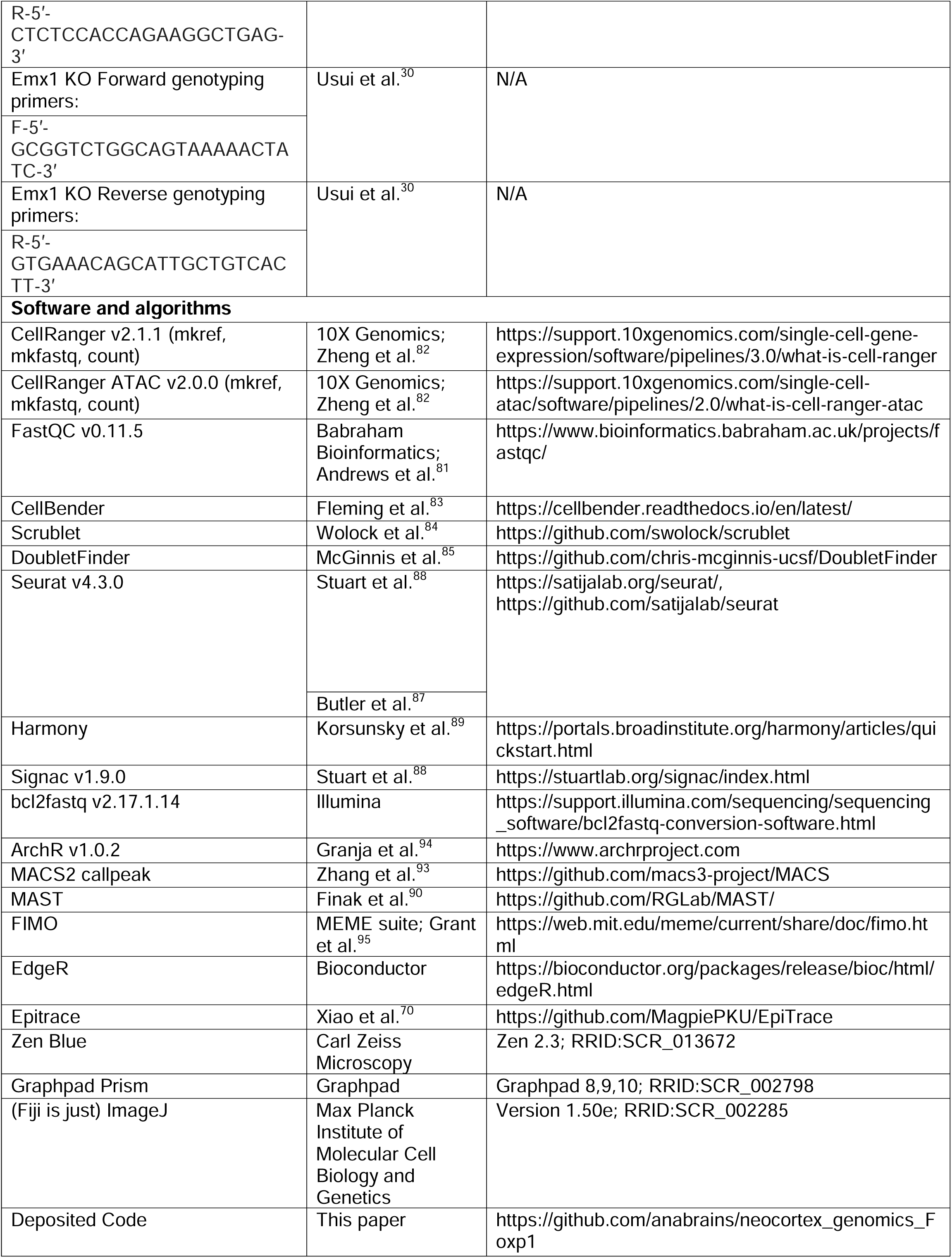

### Experimental model and subject details

#### Mice

All procedures were approved by the Institutional Animal Care and Use Committee of UT Southwestern. *Emx1-Cre ^36^* (#005628, Jackson Laboratory); *Foxp1^flox/flox^* mice ^33^, provided by Dr. Haley Tucker, were backcrossed to C57BL/6J for at least 10 generations to obtain congenic animals as previously described ^29,30^. We bred *Emx1-Cre/Foxp1^flox/flox^*males to *Foxp1^flox/flox^* female mice to obtain *Foxp1* cKO and control mice. Mice were group-housed under a 12 h light/dark cycle and given ad libitum access to food and water. Mice of both sexes were used for all experiments. For timed breeding, female mice were paired one night with male mouse and noon of the following day is considered E0.5 ^30^.

### Method details

#### Single nucleus cortical tissue processing for snRNA-seq

For embryonic timepoints, tail snips from embryos were taken for genotyping while in ice-cold PBS; then whole cortex hemispheres were rapidly dissected into separate micro-centrifuge tubes, flash frozen in liquid nitrogen, and stored at -80°C. Nuclei were isolated per modifications of other isolation protocols ^79,80^. Single nuclei were obtained from cortex samples in 1 mL ice cold Nuclei EZ lysis buffer (Sigma, #EZ PREP NUC-101), homogenized using 2 mL glass Dounce tissue grinders on ice; 15 times with pestle A and then 15 times with pestle B. The solution was then transferred to a new 2 mL microcentrifuge tube, 1 mL nuclei EZ lysis buffer was added, followed by 5-minute incubation on ice, and then centrifuged at 500xg for 5 minutes at 4°C. Supernatants were removed, pellets were resuspended in 1 mL nuclei EZ lysis buffer, followed by an additional 5-minute incubation on ice, and centrifuged at 500xg for 5 minutes at 4°C. The supernatant was then discarded, the pellet resuspended in NSB (1X PBS w/ 0.2 U/uL RNase inhibitors / 1% Ultra-pure BSA) and filtered through a 40 um Flowmi Cell Strainer (Bel-Art, H13680-0040). Nuclei were then counted on a hemocytometer (iChip) in a 10 uL of suspension with 1:1 0.4% Trypan Blue. Nuclei concentration was adjusted for targeted sequencing of 7,000-10,000 nuclei/sample using the 10X Genomics Single Cell 3′ Reagent Kits v3 protocol to prepare libraries. 4 mice/genotype and timepoint were used, including males and females. Libraries were sequenced using the Illumina NovaSeq 6000 via the McDermott Sequencing Core at UT Southwestern.

#### Single nucleus cortical tissue processing for snATAC-seq

Nuclei were isolated per modification of prior protocols ^79,80^, as described above, but using different nuclei wash buffers specialized for snATAC-Seq (10mM Tris pH 7.4, 10mM NaCl, 3mM MgCl2, 10% BSA, and 1% Tween 20). After counting, nuclei were diluted using the 20X Nuclei Buffer (10X Genomics) before proceeding with the 10X Genomic Single Cell ATAC Kit v1.1 and v2 protocol. Between 7,000 and 10,000 nuclei were targeted per sample. Three replicates per genotype for E13.5 and P0.5, and 2 replicates per genotype for E16.5, were processed for a total of 16 samples prepared in 2 batches. Libraries were sequenced using the McDermott Sequencing Core at UT Southwestern.

#### Pre-processing of snRNA-seq data

Unprocessed sequencing data were acquired from the McDermott Sequencing Core at UT Southwestern as binary base call (BCL) files. BCL files were de-multiplexed with the 10X Genomics i7 index (used during library preparation) using Illumina’s bcl2fastq v2.17.1.14 ^81^ and *mkfastq* command from 10X Genomics Cell Ranger v2.1.1 tools ^82^. Extracted paired-end fastq files were checked for read quality using FASTQC v0.11.5 ^81^. The raw gene-count matrix containing cells as rows and genes as columns was used for ambient RNA removal. Ambient RNA contamination was assessed using CellBender 0.2.0 (remove-background, default parameters), a package in Python 3.7 ^83^. The CellBender processed count matrix was used for downstream analyses.

#### Doublet removal

Scrublet ^84^ and DoubletFinder ^85^ (post-clustering) were used to predict doublets in the snRNA-seq data. Doublets identified within both packages were deemed as “true” doublets and removed from downstream analyses.

#### Intronic read ratio

As an additional measure to remove ambient RNA contamination, we used the intronic read ratio because a low intronic read ratio indicates the presence of non-nuclear transcripts ^86^. We measure intronic read ratio per cluster per sample, and clusters with an intronic read ratio below 0.5 were removed from downstream processing.

#### Clustering analysis

The described single-nuclei RNA-seq count matrix was used for clustering using the Seurat analysis pipeline in R ^87,88^. A Seurat object was created using default parameters. Then, nuclei with more than 10,000 molecules (*nUMI* per cell) and nuclei with more than 5% mitochondrial or ribosomal content, or fewer than 300 detected genes (*nFeature*) per nuclei at E13.5 or <400 at E16.5 and P0.5 were filtered out to discard potential doublets and low-quality nuclei. Genes from the mitochondrial chromosome and chromosomes X and Y were also removed as samples were from mixed sexes.

The filtered data were then log-normalized and scaled using a factor of 10,000 using *NormalizeData*, and regressed for covariates such as number of UMI per cells, percent mitochondrial content per cell, and batch, as described in the Seurat analysis pipeline ^87^. To further identify the top variable genes, the data were used to calculate principal components (PCs). Using the elbow plot and jackstraw analyses, statistically significant PCs were used to identify clusters within the data using the original Louvain algorithm. Harmony ^89^ was used to regress for batch effects using *RunHarmony*. The clusters were then visualized with uniform manifold approximation and projection (UMAP) in two dimensions using reduction = “*harmony*” ^87^.

#### Cell type annotation

Cluster marker genes were identified using *FindAllMarkers* with default parameters. Cluster markers from this study were compared with markers from published scRNA-seq datasets from embryonic and neonatal mouse cortex ^37,38^. Cell types were assigned to clusters based on statistically significant enrichment of gene sets using the hypergeometric test (with a background of the total number of expressed genes. Additionally, clusters that overlapped significantly with multiple cell types were called for the most significant overlap (smallest adj. p value) and analyzed for expression of top marker genes of known cell types. This was combined with the cell type identities assigned when using the Seurat *LabelTransfer* protocol with the reference dataset. This is the “all-cells” (*CellType1*, *CellType1_coll*) dataset. Thereafter, only cells of the excitatory neuronal lineage were selected to continue the analysis. Total cell counts were as follows E13.5: 56,557; E16.5: 36,708; P0.5: 84,075.

#### Cell type re-clustering

Cells from clusters that fell into excitatory neuronal lineage categories were taken as a subset from the “all-cells” dataset and re-clustered to define cell subtypes. Clusters that were then poorly clustered or had inhibitory gene markers were excluded from further analysis. Processing was done again as described above and cell types assigned based on canonical markers and based on statistically significant enrichment from existing datasets ^37,38^ gene sets using the hypergeometric test (with a background of the total number of expressed genes: 30,578). This dataset is referred to as excitatory neuronal lineage (*CellType*, *CellType_coll*). Total cells per timepoint are E13.5: 31,543; E16.5: 26,444; P0.5: 50,672.

#### Differentially expressed gene (DEG) analyses

To determine differentially expressed genes, pairwise differential gene expression analysis tests were performed using MAST-GLM ^90^ within each broad cell type-pair between control and cKO as previously done^91^. Thus, only one broad cluster (e.g., apical radial glia) per major cell type was used for DEG analysis (labeled *CellType_coll*). Only DEGs with an adjusted p-value < 0.05, |logFC| > 0.15 when compared to control were deemed significant ^90^. Additionally, DEGs were generated using the *CellType_coll* label with Seurat’s *FindMarkers* with test.use=*NegativeBinomial*, and with EdgeR pseudo-bulk.

#### DEG gene ontology (GO) analysis

Gene ontology (GO) analysis was performed in R using enrichGO via clusterProfiler with biological processes or molecular function ^92^. Terms were deemed significant if they had a corrected p-value < 0.05 and contained a minimum of three genes.

#### Overlap with outside gene databases

High-confidence ASD associated genes were downloaded from the SFARI-ASD ^54^ database. ASD risk genes identified by two prior studies were used ^55,56^. Synaptic genes were downloaded from Synaptome.db ^59^. Significant overlap was determined using the hypergeometric overlap test.

#### Pseudotime analysis

Pseudotime analysis was performed using destiny ^69^ in R. Data were processed as described above. For ease of visualization, within cell-type clusters (e.g., all L2-4 cells) were collapsed (*CellType_coll*), then separated by genotype per timepoint for within-timepoint, or between-genotype comparison. For each timepoint, the control and cKO object were used for creation of the diffusion map and pseudotime required for destiny. The values of pseudotime per cell and the significant difference between pseudotime estimates was tested using the Mann Whitney test using continuity correction in R.

#### snATAC-seq data Preprocessing

BCL files were processed running the *demux* command. Cell Ranger ATAC 2.0.0 was used to process Chromium Single Cell ATAC-seq data (10X Genomics), running *mkfastq*, and *cellranger-atac count* with reads aligned to mm10 reference genome (cellranger-arc-mm10-2020-A-2.0.0). Then, the peak/cell matrix was imported into Signac version 1.6.0 (https://satijalab.org/signac). Peaks were called using the Signac function *CallPeaks()*, which wraps the MACS2 function to call peaks ^93^. Doublet information was generated in ArchR ^94^ and was then exported and added to the Signac object.

Cells were retained based on the following QC metrics: peak_region_fragments > 0.02 & peak_region_fragments < 0.98, based on total sample percentages per term; pct_reads_in_peaks > 0.02; blacklist_ratio < 0.98; nucleosome_signal < 0.98; TSS.enrichment > 0.02. Common peaks were generated between biological replicates per timepoint / genotype and biological replicates merged within genotype and then between genotypes. After quality control and filtering, datasets from each of the three timepoints were analyzed as follows. Gene activities for each gene in each cell were calculated using the *GeneActivity*() function by summing the peak counts in the gene body + 2 kb upstream. Data were then normalized using term frequency inverse document frequency (TF-IDF) normalization (*RunTFIDF*), followed by dimensionality reduction using Singular Value Decomposition (*RunSVD*). K-nearest neighbors were calculated using *FindNeighbors(reduction=”lsi”, dims=2:30)*. Finally, cell clusters were identified by a shared nearest neighbor (*SNN*) modularity optimization-based clustering algorithm *FindClusters(algorithm=3, resolution=0.6)*. To correct for batch differences, harmony was called using *RunHarmony(reduction = ‘lsi’, assay.use = ‘peaks’, group.by.vars = c(’batch’, ‘orig.ident’)*. UMAP was generated using *RunUMAP* function with *reduction=“harmony*” and *dims=2:30*.

#### snRNA-seq and snATAC-seq data integration and cell-type annotations

Data from snATAC-seq were labeled based on cell labels in the corresponding snRNA-seq experiments described above at corresponding ages using Signac data integration. Briefly, cross-modality integration and label transfer was done using *FindTransferAnchors(reduction=’cca’)* and *TransferData(weight.reduction=’lsi’)* functions. Shared correlation patterns in the gene activity matrix and snRNA-seq datasets were used to match biological cell types across the two modalities. This analysis returned a classification of cell-type prediction scores for each cell. Cells were assigned the identity linked to their highest prediction score and poorly scored cells received a label of non-specific (“*Nonspec*”).

#### Cell type re-clustering

Cells annotated as part of the excitatory neuronal lineage were subset as in the snRNA-seq dataset and processed as described above for re-clustering and cell-type annotation. This subset was used as the working excitatory neuronal lineage analysis for downstream analyses, with cell types annotated under “*CellType*” and “*CellType_coll*”. This comprised 163,290 peaks and 9,111 nuclei at E13.5 from n=3/genotype; 165,957 peaks and 5,076 nuclei at E16.5 from n=2/genotype; and 188,540 peaks and 12,620 nuclei at P0.5 from n=3/genotype.

#### Differentially accessible regions

Differentially accessible regions (DARs) were calculated using the Signac package *FindMarkers* with *test.use = “LR”* and *latent.vars = c(’peak_region_fragments’,’batch’)* and compared for collapsed cell types “*CellType_coll*” of the excitatory neuronal lineage between genotypes per cell type per timepoint. Peaks for X and Y chromosomes were removed. DARs were significant if the corrected p value was less than 0.05 (adj. *p-val < 0.05*) and |logFC| > 0.25).

#### Motif enrichment

To determine overrepresented motifs in a set of differentially accessible peaks, we followed the Signac motif analysis vignette. First, motif information was added to the each timepoint dataset using *AddMotifs*(). Motif information from mm10 was added from the JASPAR2020 database in Signac. We then calculated the differential accessibility of each collapsed cell type between genotypes per timepoint, using *FindMarkers*() function with *test.use = ‘LR’* and *latent.vars = (’nCount_peaks’, ‘batch’)* in Signac. For each cell collapsed type, differential distal elements linked to genes with differential gene activity were used for motif enrichment analysis. Peaks were significant with p_val_adj <0.005 and |avg_log2FC| < 0.25. To find overrepresented motifs, we scanned a given set of differentially accessible peaks for all the DNA-binding motifs JASPAR2020_CORE_vertebrates_non-redundant databases (https://jaspar2020.genereg.net). Using *FindMotifs*(), we then computed the number of features containing the motif (*observed*) compared to the background. Background peaks were randomly sampled from all snATAC–seq peaks and matched for GC content using *MatchRegionStats()* in Signac. Enriched FOX motifs were tested for within overlapping DEG/DARs using FIMO from the MEME suite ^95^.

#### Pseudo-multiome factor analysis

For the pseudo-multiome factor analysis, we first used GLUE ^67^ to find shared embeddings for the scRNA-seq and snATAC-seq features. Next, we used the ‘GLUE_pair’ function from the OmicVerse ^96^ python package to find the nearest neighbor cell whose GLUE embeddings were highly correlated (Pearson r > 0.9) over 20 iterations. We used the paired cells to construct a multi-omics factor analysis (MOFA+) model ^68^, which we used for downstream analysis. We analyzed the model with decoupler ^97^ using a similar approach to MOFAcellular ^98^, but with composite modality-cell type views.

### BrdU or EdU experiments

To label dividing cells and measure migration, timed pregnant *Foxp1*^flox/flox^ dams were intraperitoneally injected at specified embryonic days (E12-E17) with a 50 mg/kg dose of 5-Bromo-2’-deoxyuridine (BrdU, Sigma-Aldrich B5002) per gram of body weight of a 20 mg/mL BrdU solution dissolved in PBS or with EdU (Invitrogen, C10340) at 5 micrograms per gram body weight. For migration assays, brains were collected from injection date until P7 or P16. For S-Phase experiments, EdU was injected into pregnant dams, then 1.5 hours later BrdU was injected (Ti), then embryos were collected 2 hours after the first injection (30 min post-BrdU). S-Phase length (Ts) = Ti x (EdU+BrdU+ cells/ EdU+BrdU-cells); and cell cycle length (Tc) was calculated by Ts / (EdU+BrdU+ cells/ Ki67+ cells) as Ki67 marks all cycling progenitors ^71^. All BrdU treated brains underwent antigen retrieval. For migration assays, sections were stained with anti-BrdU and layer-specific antibody markers. For EdU treated brain, Click-iT EdU kit instructions were followed (Invitrogen, C10340). Images were acquired at 20X or 63X using a Zeiss LSM 880 confocal microscope. Stitched maximum intensity projection images are used for cell counting using FIJI.

### Immunohistochemistry

For embryonic timepoints, dams were anesthetized with CO_2_ and embryos were extracted rapidly and placed in a Petri dish with ice-cold 1X PBS. Tail snips were taken for genotyping. Embryos we/re transferred to clean Petri dishes with ice-cold 1X PBS twice before drop fixing in 4% PFA at 4°C for 30 min-overnight. Embryos were cryoprotected in 20% and 30% sucrose with 0.01% sodium azide until they sunk before mounting in OCT media and freezing on dry ice. Sections were made at a thickness of 12-30 um on a Leica CM1950 Cryostat, directly mounted onto slides, and allowed to dry before processing. Neonatal and adult mice were anesthetized then transcardially perfused with 1x PBS and 4% PFA and their brains postfixed overnight in 4% PFA at 4°C. After cryoprotection in 20% and 30% sucrose with 0.01% sodium azide overnight at 4°C, brains were embedded in Tissue-Tek CRYO-OCT Compound and cryosectioned at 15-30 um.

For staining, all washes were done with TBS or 0.4% Triton X-100 in TBS (TBS-T) unless otherwise stated. When specified, antigen retrieval was performed in citrate buffer (10 mM tri-sodium citrate, 0.05% Tween-20, pH 6) for 10 min at 95°C. Free aldehydes were quenched with 0.3 M glycine in TBS-T for 1 hour at room temperature (RT), and sections blocked in 10% normal donkey serum (NDS) and 3% bovine serum albumin (BSA) in TBS. Sections were incubated 1.5 hours at RT or overnight at 4°C in primary antibodies diluted in 10% NDS and 3% BSA in TBS. Secondary antibody incubations were performed for 1 hour at RT or overnight at 4°C in 10% BSA in TBS. Sections were then washed three times with TBS. Sections were incubated with Hoechst or DAPI 1:2000 for 2 min, then washed in TBS. Coverslips were mounted using ProLong Diamond Antifade Mountant without DAPI (#P36970, Thermo Fisher).

### Imaging and image analysis

Images were acquired using a Zeiss LSM 880 confocal microscope at the UT Southwestern Neuroscience Microscopy Facility and processed and analyzed using Zeiss ZEN Lite and FIJI. For quantifications, tile scan Z-stack images of the region of interest were acquired at 20X magnification from similar coronal sections across at least 2–3 mice/genotype across at least 2 mouse litters. Stitched maximum intensity projection images are used. Cell counting was done using the FIJI Cell Counter plugin. Layer thickness was measured in FIJI and normalized to total cortical thickness based on DNA stain DAPI or Hoechst. Experimenters were blind to genotypes during quantification.

### Migration assays quantification

Pregnant dams were injected with EdU (5 ug/g) at embryonic date of interest to label cells in S-Phase for migration. Offspring were then collected postnatally at P7.5 or P16.5. Antibodies for EdU, CUX1, and CTIP2 were used to denote cells proliferating at the time of injection, and markers of UL and DL cells respectively. Location in cortical layers for cells co-positive for EdU+CUX1+ were binned for UL and DL based anatomically on marker genes CUX1, CTIP2, and Hoechst staining. Specifically, to denote migration defects, EdU+CUX1+ cells in L5-6 were measured. Then overall locations of EdU+ cells in cortical layers were semi-automatically quantified using FIJI and grouped into UL or DL bins and compared using the Mann-Whitney test.

### Cell cycle exit assay quantification

Pregnant dams were IP injected with EdU and brains from embryos from the pregnant dams were collected 24 hours later. Co-staining of coronal sections 12-25 um thick with Ki-67 and EdU was used to assess the state of cells 24 hours post injection, e.g., whether they remained proliferative, KI67+ (Ki67 is a proliferating cell marker that is expressed in all mitotic cells). Cells that were EdU+Ki67- at the time of collection were cells that exited the cell cycle, while EdU+Ki67+ cells were still in the cell cycle; these were both normalized to total EdU cells. Analyses were done in FIJI and Mann-Whitney test done in GraphPad Prism.

### Single molecule fluorescent in situ hybridization (smFISH) and quantification

Brains were collected from P1 mice, embedded in OCT compound and snap frozen in isopentane pre-cooled with liquid nitrogen. Frozen brains were sectioned at -20°C into 15 µm thick coronal slices targeting the somatosensory cortex. The sections were thaw-mounted onto Superfrost Plus Microscope slides (*ThermoFisher Scientific, cat #12-550-15*). Single molecule Fluorescent in situ hybridization (smFISH) was performed using the RNAscope Multiplex Fluorescent Reagent Kit v2 assay (*Advanced Cell Diagnostics, cat# 323100*) as per the manufacturer’s protocol for fresh frozen tissue, except that slides were baked for 30 minutes at 60°C before fixation and Protease III treatment was applied for 15 minutes at 40°C.

RNAscope probes for *Ctip2* (*also known as Bcl11b, ACD, cat# 413271-C2 and C3*), *Rora* (*ACD, cat# 520031-C1*)*, Kirrel3* (*ACD, cat# 463651-C1*) and *Nrxn3* (*ACD, cat# 505431-C3*) were used to identify the expression. The probes were incubated with the tissue to hybridize with their respective target mRNAs. Opal fluorophores 520 (*FP1487001KT, Akoya Biosciences, 1:750*), 570 (*FP1488001KT, Akoya Biosciences, 1:750*) and 690 (*FP1497001KT, Akoya Biosciences, 1:750*) were used to label the gene-specific probes after signal amplification.

Images were acquired using a Zeiss LSM 880 confocal microscope at 20x magnification at the UT Southwestern Neuroscience Microscopy Facility. One image per section was captured from the somatosensory Four square-shaped bins were sampled from each image to encompass the entire *Ctip2+* layer (Layers 5-6). The total number of pixels corresponding to each gene of interest was quantified using custom ImageJ Macro code and R script in Fiji and R respectively. The total pixels were normalized with DAPI+ cell counts. Statistical analysis was performed using a linear mixed model with genotype as the fixed factor and individual as the random factor, nested with sections, hemisphere and bins. A total of three mice per genotype were analyzed, with 2-3 sections from each mouse.

### Cell type proportion calculations

To calculate differences in cell type proportions between genotypes, we take the number of cells in each broad cell type group (*CellType_coll*) over the total cell types per genotype and compare between genotypes using the two-proportions z-test with Yates correction which performs continuity correction for large values. For each time-point of the snRNA-seq dataset, there are 3 replicates per genotype.

### EpiTrace mitotic age inference

We used EpiTrace ^70^ v0.0.1.3 in R to conduct snATAC-based mitotic age inference. Briefly, EpiTrace infers cellular mitotic age by counting the fraction of DNA methylation-age-associated genomic loci with open chromatin in snATAC-seq data and compares it to a reference. We used the reference derived from the Illumina MM285 methylation array and mapped to the mm10 reference genome. We ran EpiTrace separately on Seurat (v 5.1.0) objects from each experimental timepoint with nuclei from both the control and knockout.

### Quantification and statistical analysis

Statistical methods and code used for snRNA-seq and snATAC-seq analysis are provided in the methods section. Two-three animals per genotype at each time point were used. DEGs were determined using EdgeR and considered significant if adj. p-val < 0.05 and average |logFC| > 0.15 when compared to control samples. DARs were determined in R using Signac by comparing knockout against control genotypes using the *FindMarkers* command and likelihood ratio test (LR) with *latent.vars=’peak_region_fragments’, ‘batch’*. DARs were considered significant if p_val_adj <0.05 and |avg_log2FC| < 0.25. Peaks were compared across a pairwise comparison and used as background for motif analysis using *MatchRegionStats*. Tests for differences in proportion were done in R using the two-proportions z-test with Yates correction. All statistical tests for cell cycle, IHC, and smFISH were performed using GraphPad Prism to obtain p values (Mann Whitney test) and test normality. Statistical tests used for each analysis are described in depth in figure legends. Sample sizes are indicated in figure legends. Sample size represents number of animals in IHC, snRNA-seq, and snATAC-seq analysis. Sample size represents the number of cells in smFISH analysis. All graphs are displayed as mean ± SEM.

---

**Table S1. DEGs and DARs for each broad cell type, overlapping E16.5 DARs and E15.5 ChIP-seq peaks, and overlapping PO DARs and E18.5 ChIP-seq peaks, related to Figures 1 and 2**.

**Table S2. Enriched motifs per age and cell type, related to Figure 2**.

**Table S3. Gene ontology for Biological Process or Molecular Function for DEGs and DARs, related to Figure 4**.

